# Structure-guided point mutations on FusionRed produce a brighter red fluorescent protein

**DOI:** 10.1101/2020.04.20.051763

**Authors:** Srijit Mukherjee, Sheng-Ting Hung, Nancy Douglas, Premashis Manna, Connor Thomas, Annika Ekrem, Amy E. Palmer, Ralph Jimenez

## Abstract

The development of fluorescent proteins (FPs) has revolutionized biological imaging. FusionRed, a monomeric red FP (RFP), is known for its low cytotoxicity and appropriate localization of target fusion proteins in mammalian cells but is limited in application by low fluorescence brightness. We report a brighter variant of FusionRed, FusionRed-MQV, which exhibits an extended fluorescence lifetime (2.8 ns), enhanced quantum yield (0.53), higher extinction coefficient (~140,000 M^−1^cm^−1^), increased radiative rate constant and reduced non-radiative rate constant with respect to its precursor. The properties of FusionRed-MQV derive from three mutations - M42Q, C159V and the previously identified L175M. A structure-guided approach was used to identify and mutate candidate residues around the phenol and the acylimine ends of the chromophore. The C159V mutation was identified via lifetime-based flow cytometry screening of a library in which multiple residues adjacent to the phenol end of the chromophore were mutated. The M42Q mutation is located near the acylimine end of the chromophore and was discovered using site-directed mutagenesis guided by x-ray crystal structures. FusionRed-MQV exhibits 3.4-fold higher molecular brightness and a 5-fold increase in the cellular brightness in HeLa cells (based on FACS) compared to FusionRed. It also retains the low cytotoxicity and high-fidelity localization of FusionRed, as demonstrated through assays in mammalian cells.

## Introduction

The availability of genetically-encoded fluorophores such as fluorescent proteins (FPs) initiated a revolution in biological imaging.^1^ Routine use of FPs in assays involving technologies such as Förster resonance energy transfer (FRET),^2^ fluorescence lifetime imaging microscopy (FLIM),^3,4^ molecular sensing, ^5^ and nanoscopy, ^6^ make them indispensible tools for biological research. Current efforts focus on developing FPs with excitation and emission in the far-red/near-infrared wavelengths and with photophysical properties such as photoswitching and fluorescence intermittency optimized for super-resolution imaging modalities.^7–15^ Nevertheless, all imaging applications benefit from increased cellular brightness, which is strongly dependent on molecular brightness (defined as the product of the molar extinction coefficient (**ε**) and the fluorescence quantum yield (**ϕ**)).^16 17^ Lifetime-based selection methods have exploited a correlation between fluorescence lifetime (**τ**) and **ϕ** to develop FPs with higher **ϕ** such as NowGFP,^18^ mTurquoise^2,19^ and mScarlet, which is the brightest red FP.^20^

FPs selected for higher **τ** generally show a decreased rate constant of non-radiative decay (k_non-rad_), ultimately resulting in increased **ϕ** and higher brightness. However, **ϕ** is also linearly related with k_rad_, therefore there is an interplay between the absorption probability and radiative emission probability. The Strickler–Berg equation formalizes the relationship between the k_rad_ and **ε** as well as other spectral properties such as the energies and profiles of the absorption and emission bands.^21^ For example, blue shifts and decreased spectral width can lead to higher probabilities of radiative decay.^22^ The relationship between spectral characteristics and the rate constants of energy decay have recently been discussed for FPs.^23, 24^ An engineering strategy that maximizes the radiative rate constant while simultaneously minimizing the non-radiative rate constant should be an effective way to brighten a FP.

These issues are particularly acute for red FPs (RFPs), which generally have lower values of **τ** and **ϕ** compared to shorter wavelength variants, and are therefore not as bright. ^25^ The RFP chromophores contain an acylimine moiety, which expands their electronic conjugation and leads to a ~50 nm red shift in their absorption and emission spectra.^26,27^ Many potential non-radiative decay mechanisms that lead to lower quantum efficiencies in such systems have been investigated, including transitions to dark states; ^28, 29^ charge accumulation and twisting of the acylimine moiety; ^30^ changes in hydrogen bonding patterns; and electrostatic, steric and conformational effects associated with their increased number of vibrational degrees of freedom.^31–33^

Cellular brightness depends on both photophysical and non-photophysical factors such as translational efficiency, protein folding and chromophore maturation.^34^ FPs like FusionRed-M (FR-M) and mScarlet-I were developed to enhance cellular brightness over their molecularly brighter parents FR-1 and mScarlet, respectively.^20,35^ Although there have been many efforts to improve the cellular and molecular brightness of FPs, performance in cell biology applications may still suffer from cellular toxicity, poor or over expression in certain cellular contexts and *in cellulo* oligomerization.^16,34^

FusionRed (FR) was developed as a non-cytotoxic RFP that shows efficient and appropriate localization in multiple fusion constructs in different organisms.^36^ However, this FP has not been widely adopted because it is relatively dim, in part due to its low **ϕ** (0.19). We recently developed the FR-M variant (FR-L177M, numbered with respect to mCherry and FR-L175M numbered with respect to FR; See Supplementary Information S1: Table S1), which is 1.9-fold brighter than FR in HeLa cells.^35^ In this study we use structure-guided engineering strategies to identify substitutions M42Q and C159V (sequence numbering with respect to FR) facing into the beta barrel that individually increase the **τ** of FR. These mutations lead to the development of the bright triple mutant FR-MQV. We perform detailed *in vitro* photophysical analysis which corroborate the increase in the molecular brightness in FR-MQV based on minimizing k_non-rad_ and increasing the k_rad_. Additionally, we investigate the cellular properties of FR-MQV and find it is 5-fold brighter in HeLa cells (using FACS) compared to the parent FR, while preserving its physiological properties such as low cytotoxicity and high-fidelity localization.

## Results

### Lifetime-based selection of a site-directed library at the phenol end of the chromophore identifies the C159V mutation

Four positions near the phenol end of the chromophore—C159, M161, V196 and H198—were simultaneously mutated in FR to generate a ~7,000-member library (See Methods and Materials for details of library generation). We employed lifetime-based microfluidic flow cytometry^35^ to select variants with increased brightness and values of **τ** different from the parent (Figure 1). Sequencing of selected clones revealed that H198 and V196 were conserved but positions 159 and 161 showed sequence diversity (Supplementary Information S2: Table S2). Furthermore, the selected population contained two clones, FR-C159V and FR-C159L, with higher cellular brightness than the parent FR, but with differing lifetimes. FR-C159V had an increased lifetime of 2.1 ns, whereas FR-C159L had a decreased lifetime of 1.2 ns (c.f. 1.8 ns lifetime for FR: Details in Supplementary Information S2: Table S2). FR-159V was brighter and had a longer lifetime in comparison to FR, and therefore was selected for combination with the mutations identified below.

**Figure 1:**
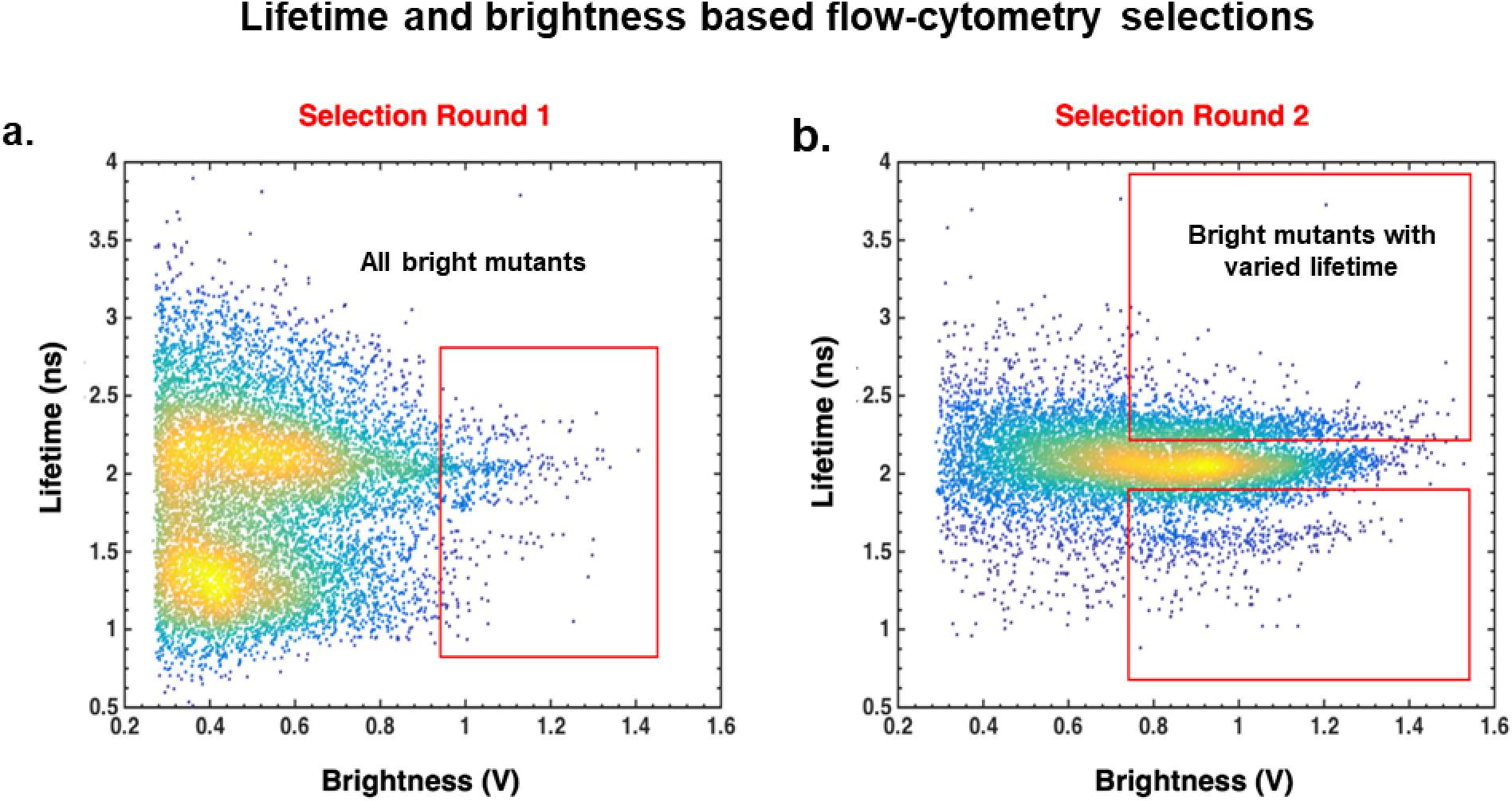
The fluorescence lifetime (vertical axis) vs. brightness (horizontal axis) screening dot-plots of the FR site-directed library targeting the phenol end, after two rounds of fluorescence-activated cell sorting (FACS) enrichments. Pseudocoloring represents normalized cell counts, where yellow indicates higher cell count and blue indicates a lower cell count. Red boxes indicate the selection gates for multi-parameter microfluidic sorts. (a) The first round of sorting for selecting bright clones. (b) The second round of sorting was performed in two batches using sorting gates for high brightness with different lifetime ranges. The mean fluorescence lifetime of FR measured on this instrument was 2.05 ns.

### The M42Q mutation on the acylimine end of FR increases the max, ε_max_Φ and τ

Mutations that alter the hydrogen-bonding structure surrounding and interacting with the acylimine end of the chromophore produce RFPs with extended Stokes shifts.^37–40^ In addition to the site-directed library aimed at mutating the phenol end of the chromophore, we investigated the acylimine end of the chromophore in the crystal structures of mKate (PDB ID:3BXB), the far-red emitting FP TagRFP-675 (PDB ID:4KGE) and FR (PDB ID:6U1A) using PyMol ^41^ (Figure 2 and Supplementary Information S3: Figure S1). Based on the structural similarity, conformational freedom and the fact that the sidechain at position 42 is a known hot-spot for altering the hydrogen-bonded chemistry in mKate and TagRFP-675,^38^ we incorporated a single point mutation M42Q into FR. The FR-M42Q (or FR-Q) mutant shows a 41% increase in **max**, an 18% increase in **ε**_**max**_ and a 42% increase in **ϕ** (Table 1 and Figure 3). The mutation also resulted in blue-shifted and narrower absorption and emission spectra and a minor change in Stokes shift (Figure 3a and Supplementary Information S4a: Figure S2, S4b: Table S3, and S4c: Figure S3). These properties reflect the higher radiative rate constant calculated for this mutant in comparison to FR (Figure 3d). Consequently, we performed site-saturation mutagenesis (library size = 20) at this position in the context of FR-M and found that the parent, along with FR-M M42Q (FR-MQ) and FR-M M42I (FR-MI) were the only variants with observable red fluorescence. FR-MQ was the brightest species with a **ϕ** of ~43% and a further increase of **τ** by 0.3 ns. FR-MI showed decreases in **ε**_**max**_ and **ϕ**, thus we did not pursue engineering of this variant. We investigated the effects of analogous mutations on closely related RFPs and noted rather different outcomes. For example, mKate M42Q shows a decrease in the molecular brightness due to ~50% decreases in both **max** and, along with a blue-shifted absorption spectrum and a significantly increased Stokes shift, which was consistent with previous findings.^38^ In addition, analogous mutations at position A44 on mCherry and mScarlet-I RFPs resulted in non-fluorescent clones.

**Table 1:** Summary of *in vitro* photophysical properties of FR mutants developed in this study vs. re-measured values for some well-characterized RFPs. Error bars have been reported measurements where independent triplicates were performed. The mCherry and mScarlet, were used as references for the **ϕ** measurements. ^16,20^

| FP | $\lambda_{\text{abs}}$<br>(nm) | $\lambda_{\text{em}}$<br>(nm) | $\tau$<br>(ns) | $\phi$ | $\epsilon_{\text{max}}$<br>(M<sup>-1</sup>cm<sup>-1</sup>) | Molecular<br>brightness | $k_{\text{rad}}$<br>($\mu\text{s}^{-1}$) | $k_{\text{NR}}$<br>($\mu\text{s}^{-1}$) |
| --- | --- | --- | --- | --- | --- | --- | --- | --- |
| FR | 574 | 596 | 1.78±0.04 | 0.24±0.04 | 94000±7000 | 1 | 135±23 | 427±26 |
| FR-M | 571 | 591 | 2.13±0.05 | 0.34±0.02 | 78000±10000 | 1.2 | 160±10 | 310±15 |
| FR-Q | 568 | 587 | 2.10±0.05 | 0.33±0.01 | 133000±2000 | 2 | 157±6 | 319±13 |
| FR-MI | 575 | 595 | 1.75±0.02 | 0.26±0.04 | 82000 | 1 | 149±12 | 423±13 |
| FR-MQ | 567 | 586 | 2.45±0.08 | 0.43±0.03 | 135000±2000 | 2.7 | 176±14 | 233±19 |
| FR-MQV | 566 | 585 | 2.77±0.07 | 0.53±0.03 | 144000±3000 | 3.4 | 191±12 | 170±16 |
| mScarlet-I | 570 | 591 | 3.26±0.07 | 0.59±0.03 | 102500±4500 | 2.7 | 184±10 | 128±12 |
| mKate | 588 | 634 | 2.26±0.07 | 0.33 | 48000 | 0.7 | 130 | 264 |
| mKate-M42Q | 577 | 654 | 2.16±0.05 | 0.17 | 25000 | 0.2 | 79 | 384 |
| mCherry | 586 | 607 | 1.67±0.07 | 0.22 (ref) | 75000±5000 | 0.8 | 137 | 488 |
| mScarlet | 569 | 592 | 3.87±0.07 | 0.71 (ref) | 103000±4500 | 3.3 | 186 | 72 |

**Figure 2:**
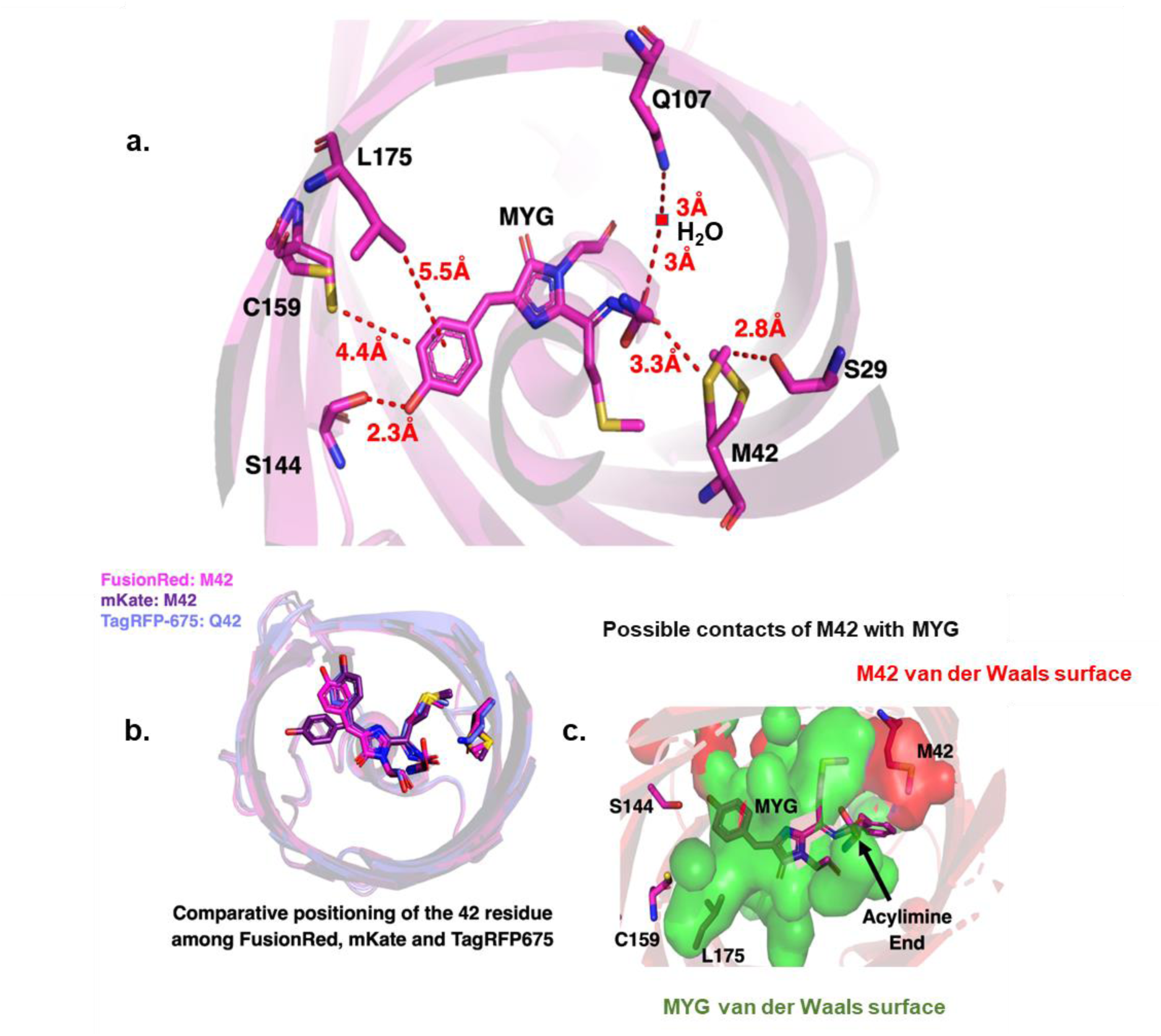
(a) Crystal structure of FR with positions around the chromophore relevant to the study generated using PyMol. (b) 42 position in TagRFP-675 (PDB ID: 4KGE, blue, 42Q), mKate (PDB ID: 3BXB, purple, 42M) and FR ?(PDB ID:6U1A, pink, 42M). The crystal structure of mKate (PDB ID: 3BXB) indicates that the phenolate end of the chromophore exist in both cis and trans forms, with the cis form overlapping with that of TagRFP-675. For FR, the fluorescent chromophore exists in the cis form as well. (c) The Van der Waals contact surfaces for the chromophore (green) versus the occupancy for the M42 residue in the FR crystal structure.

**Figure 3:**
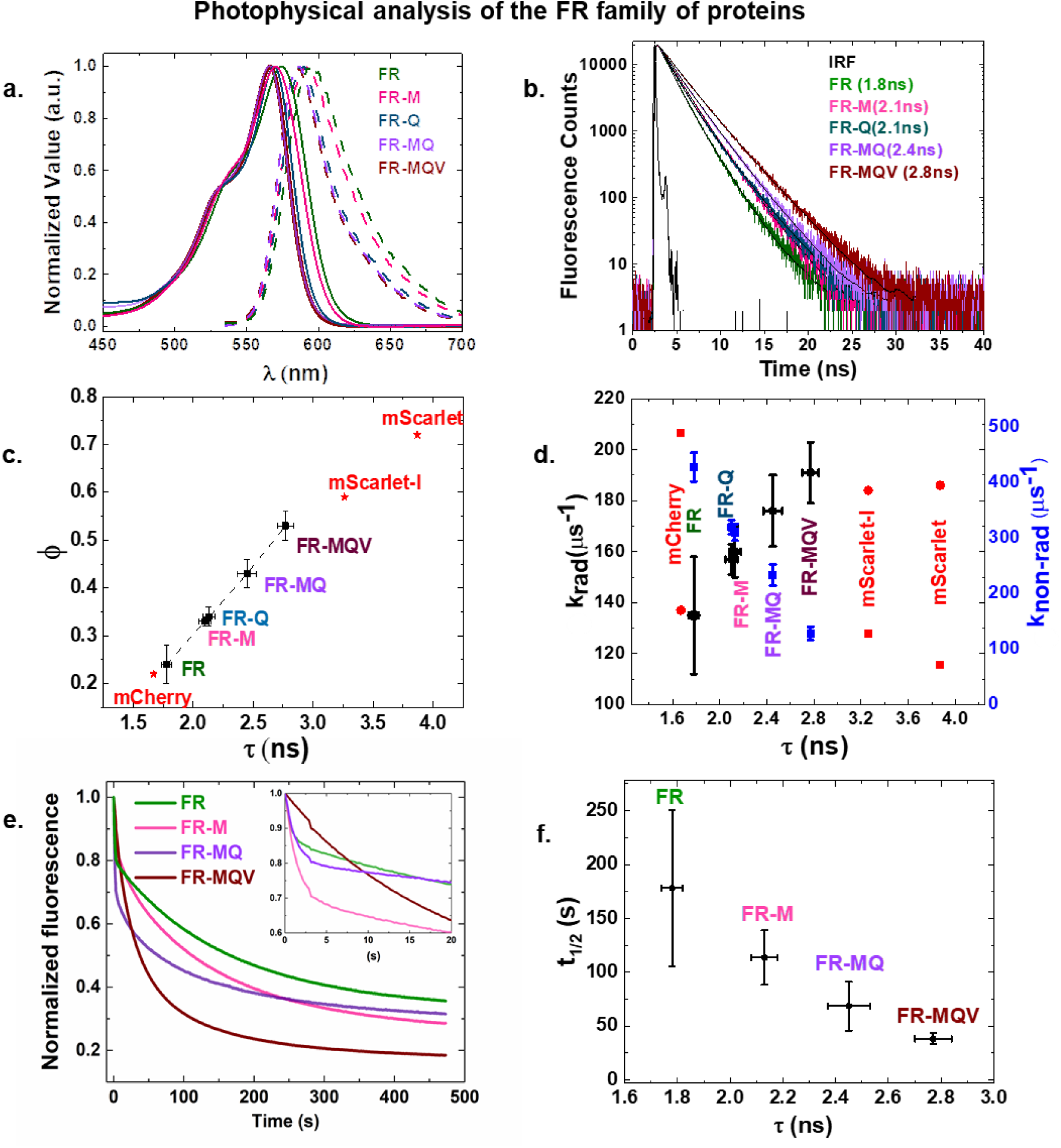
Photophysical properties of the FR family of FPs (a) Absorption (solid lines) and emission (dashed lines) spectra. (b) The TCSPC lifetime traces. Longer decay profiles indicate the gradual increase in fluorescence lifetime. The black lines indicate exponential fitting for the decay profiles. (See Supplementary Information S4g: Table S4 for fitting details) (c) Plot of quantum yield (**ϕ**) versus lifetime (**τ**). The dashed line indicates the linear rise of **ϕ** with increments in **ϕ** for the FR family of FPs. mCherry, mScarlet and mScarlet-I have been reported in red based on literature values. ^16,20^ (d) A plot showing the increase in radiative rate constant (black) and decrease in non-radiative rate constant (blue) across the FR family relative to mCherry and mScarlet (red). Vertical error bars (blue and black) in the FR family indicate the uncertainty in the calculated rate constants, the horizontal error bars (black) are the standard deviation error in the measurement of **τ**. (e) Photobleaching traces for the family of FR mutants, determined by averaging ~10 decay traces in *E. coli* cells for each FP under normalized excitation rates. (f) The dependence of the fluorescence decay half-life on the excited state lifetime.

In the course of measuring the **ε**_**max**_ with the alkali-denaturation method, we observed that M42Q containing mutants of FR produce a single hydrolysis product with an absorption band centered at 380 nm (Supplementary Information S4d: Figure S4). Most RFPs, such as mScarlet, mCherry and mRuby3, display GFP-like degradation with an absorption band at 450 nm. The hydrolysis products of FR variants with the native 42M have both 380 nm and 450 nm absorption peaks. The 380 nm band is assigned to the cleavage of the chromophore from the β-carbon of the Tyr moiety in the chromophore.^42–44^ The absence of the 450nm band in FR-Q and FR-MQ reveals alteration of the chromophore hydrolysis chemistry. This suggests that the 42Q residue interacts strongly with the chromophore in FR.

### Combining mutations M42Q, C159V and L175M led to the generation of molecularly bright FR-MQV

Each individual mutation results in an increment of **τ** by ~0.3 ns in FR. Therefore, we incorporated all three mutations into FR to yield FR-MQV. The FR-MQV mutant exhibits a **τ** of 2.8 ns which results in a high **ϕ** of 53% (Figures 3b and 3c). The absorption and emission spectra are similar, with a slight blue shift and narrowing of the spectra in all variants possessing the M42Q mutation (Figure 3a and Supplementary Information S4b: Table S3). The molecular brightness of FR-MQV is ~3.4-fold larger than that of FR. Overall, we see increases ina**τ** and **ϕ**, as a result of an increase in the radiative rate constant and a concomitant decrease in the non-radiative rate constant (Figure 3d). Fluorescence excitation spectra and wavelength-dependent lifetime measurements were collected to assess the heterogeneity of chromophore formation. We found that the FR-MQV chromophore matures predominantly to a single red-emitting spectral form with the absence of a green-emitting chromophore (Supplementary Information S4e: Figure S5; S4f: Figure S6; and S4g: Table S4). FR-MQV has a low in vitro pKa of ~4.6, like other members of the FR family (Supplementary Information S4h: Figure S7 and Table S5) and breaks down into a single product of alkali hydrolysis, resembling FR-Q and FR-MQ (Supplementary Information: S4d). We also found that FR-MQV was ~2-fold brighter than FR in E. coli (Supplementary Information S4i: Figure S8).

The photobleaching traces of FR-MQV and related variants are shown in Figure 3e. We found that under excitation-normalized conditions, longer **τ** is correlated with faster photobleaching (Figure 3f). Furthermore, it was observed that under continuous irradiance of ~5 W/cm^2^ at 560 nm, the fluorescence kinetics of FR variants with a C159 residue show two decay timescales. This behavior is typical of most FPs, which show an initial decay due to dark state conversion (also sometimes called reversible photobleaching), along with a slower timescale of permanent photobleaching.^45,46^ The C159V and C159L mutations, which incorporate aliphatic groups, significantly reduce the amplitude of the faster decay component (Supplementary Information S4j: Figure S9a). To verify that the faster component corresponds to a reversible process, we used alternating pulses of 560 nm and 438 nm excitation (Supplementary Information S4j: Figure S9b and c). The FR-C159V variant showed little recovery and FR-C159L showed slight recovery of the fluorescence, but the recovery was much larger for FR (~20% higher). These results show that the reversible photobleaching dynamics of FR are influenced by the chemical nature of the sidechain at position 159.

### FR-MQV is 5-fold brighter in HeLa cells, retains the physiological and high-fidelity localization properties of FR

Flow cytometry and confocal microscopy were employed to quantify the cellular brightness of FR-MQV in HeLa cells (Figure 4). FACS screening of HeLa cells expressing FPs fused to histone H2B protein after 48 hours of transfection revealed that FR-MQV is 5-fold brighter than the parent FR, and ~1.4-fold less bright than mScarlet (Figure 4a and Supplementary Information: S5a Table S6). Measurements with confocal microscopy on the same construct revealed that FR-MQV is ~2.3-fold brighter than FR and ~1.4-fold dimmer than mScarlet (Figure 4b and Supplementary Information: S5a Table S7). We next investigated whether the mutations in FR-MQV would influence cytotoxicity, chromophore maturation and localization of fusion proteins. FR-MQV exhibited low cytotoxicity, like the parent FR and contrary to mCherry, which showed ~80% relative decrease in cells expressing the FP after 6 days of expression (Details of the assay are discussed in Supplementary Information S5b: Figure S10). FR-MQV showed chromophore maturation kinetics identical to the parent FR (t_1/2_ ~195 min at 37 °C; Supplementary Information S5c: Figure S11 and Table S8) and similar to mScarlet (t_1/2_ ~ 130 min at 37 °C), but slower than mScarlet-I (t1/2 ~45 min at 37 °C).

**Figure 4:**
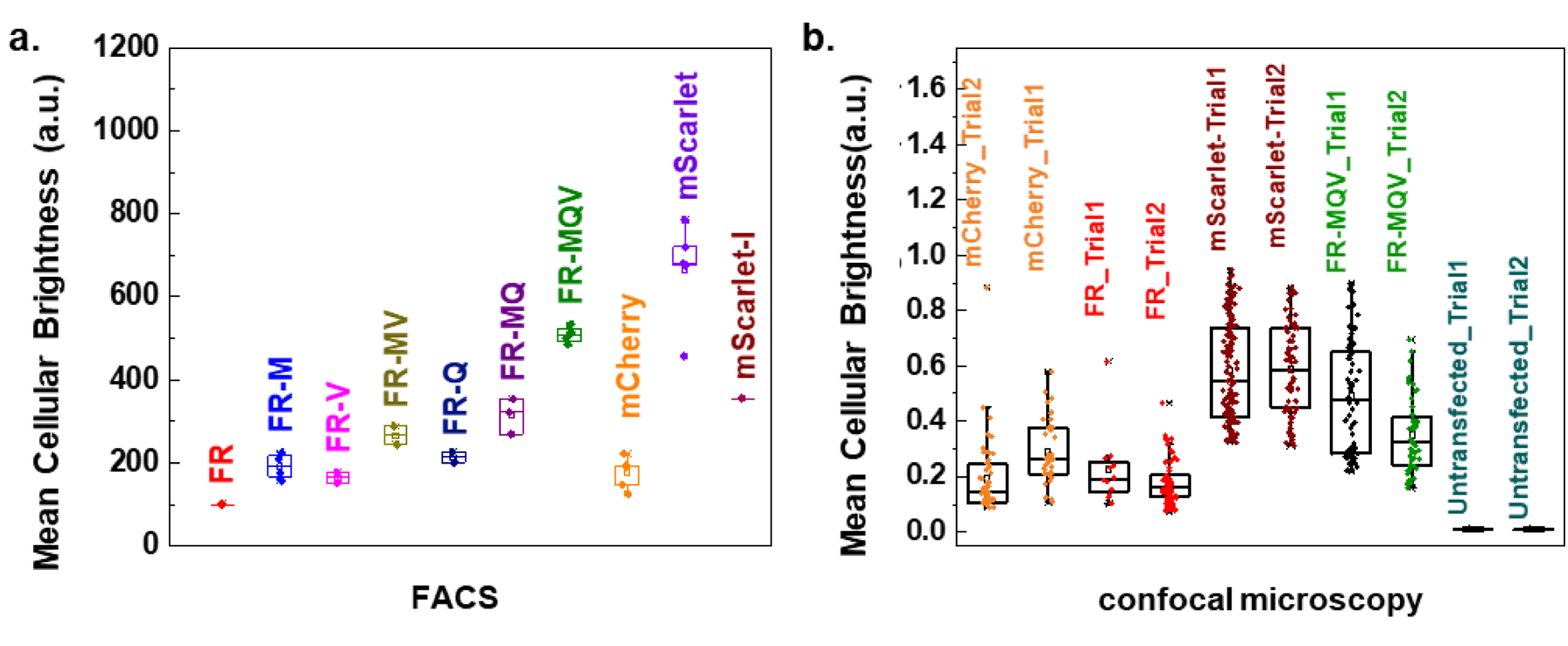
(a) FACS-based brightness assays of FPs expressed as histone H2B fusions in HeLa cells. Each biological replicate involved three technical replicate screens of ~10,000 individual cells. (b) Mean brightness values of individual HeLa cells obtained from confocal microscopy for histone H2B fusions. Each dish was an independent biological replicate and contained ~100 cells. For both a and b, the cells were sampled ~48 hours post transfection. Details of the assays are provided in Supplementary Information S5a.

Over-expression and oligomerization in cells can lead to unwanted effects such as self-quenching, mis-folding, mis-localization and disruption of organelle structure and function.^47,48^ Hence, we performed the organized smooth endoplasmic reticulum (OSER) ^49^ assay and examined localization of a GalT-FP fusion to assess the *in cellulo* monomericity and localization properties of FR-MQV relative to FR and the other bright RFPs mScarlet and mScarlet-I.

In the OSER assay, FPs are fused to the cytoplasmic end of the endoplasmic reticular signal anchor protein (cytERM).^49^ Membrane localization increases the local concentration leading to the formation of dimers or higher order oligomers that distort the ER structure which are observed as cellular sub-structures or “whorls”. Figure 5a shows confocal microscopy images of cytERM-FPs expressed in U2OS cells and the corresponding OSER scores. TagRFP-T (positive control) shows characteristic oligomerization in the form of sub-cellular “whorls”, while the FR mutants, mScarlet and mScarlet-I generally lack such structures (See Methods and Materials for details of experiments and data analysis). In this work, we measured a score of 88 for FR and 87 for FR-MQV which are ~5% higher than measured for mScarlet (84) and ~10% higher than mScarlet-I (80). We also developed GalT-FP fusions to localize the FPs of interest to the Golgi. Proper and mis-localization to the Golgi in U2OS cells imaged using confocal microscopy are shown in Figure 5b. Our analysis indicates that FR-MQV and other FR-derived mutants outscore mScarlet and mCherry in terms of proper localization to the Golgi. Our results are qualitatively consistent with previous observations where FR correctly localizes to the Golgi, but GalT-mCherry frequently displays puncta in the cytoplasm.^50^ We observed higher occurrences of mis-localization in the bright RFPs mScarlet and mScarlet-I compared to FR mutants. Thus, cellular imaging studies indicate that FR-MQV retains the high-fidelity localization properties of FR without compromising low cytotoxicity and appropriate maturation and expression.

**Figure 5:**
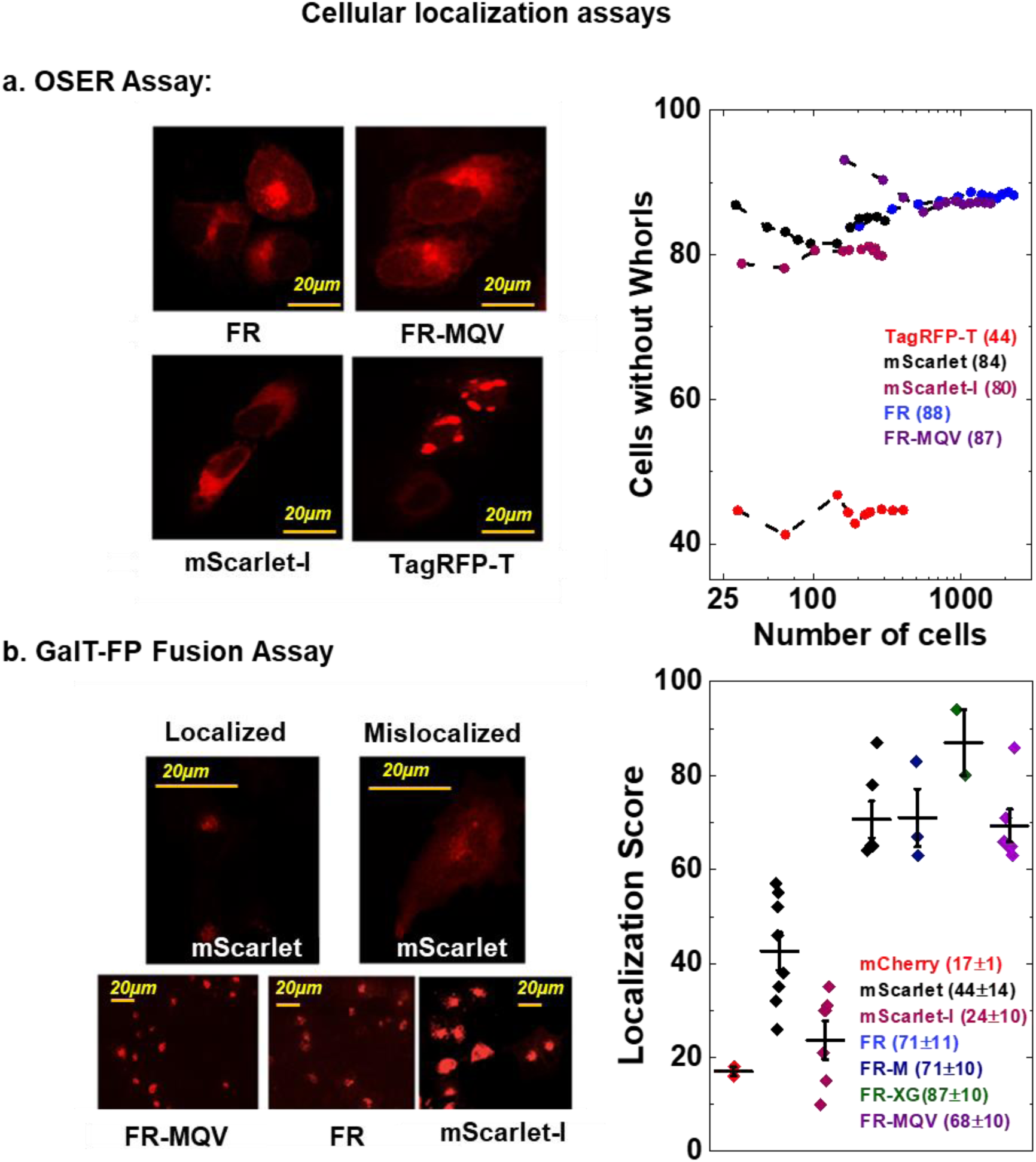
(a) *The OSER Assay:* Representative maximum intensity projected confocal images for U2OS cells expressing cytERM-FP constructs for FR, FR-MQV, mScarlet-I and TagRFP-T (negative control), and the graphical representation of the OSER scores for these cells. The y-axis on the plot indicates the percent cells that do not express sub-cellular “whorls”. The OSER scores for this assay were: FR, 88 (2,250 cells), FR-MQV, 87 (1,580 cells), mScarlet, 84 (306 cells), mScarlet-I, 80 (291 cells) and TagRFP-T, 45 (406 cells). (b) *The GalT-FP Fusion Assay:* Confocal microscope images for U2OS cells expressing the GalT-FP fusion constructs for mScarlet, FR-MQV, FR and mScarlet-I, with the localization scores (indicative of the percent cells with the FP localized correctly to the Golgi) plotted for each RFP. The microscopy images show cells expressing GalT-mScarlet localized and mis-localized to the Golgi.

## Discussion

In this work we engineered FR using a structure guided approach, aimed to develop mutants with increased **τ** manifesting in higher **ϕ**, eventually leading to a FP with higher molecular and cellular brightness. We examined both the phenol and the acylimine ends of the chromophore for sites where we could potentially improve steric packing, increase rigidity, interact chemically or electrostatically with the chromophore. At the phenol end, this approach led us to mutate multiple residues, thus, the need for lifetime screening of a 7,000-member library. In contrast, at the acylimine end we identified the capacity for an interaction to be added at position M42. Crystal structure data suggests conformational freedom of the sidechain at position 42 in FR.^51^ The position 42 lies in close proximity to the acylimine end of the chromophore and based on our previous work, is critical to the hydrogen-bond network on that end of the chromophore in closely related FPs mKate and TagRFP-675.^38^ A smaller site-saturated library of 20 possible mutants at position 42 allowed us to carry out a plate based screen on FR-M, where Gln stood out as the best possible sidechain residue in terms of increases in **τ** and **ϕ**.

Individually L175M, C159V and M42Q substitutions, increased the **τ** of FR by ~0.3 ns (~17% increase), and when incorporated together in FR-MQV exhibited ~1 ns (~55%) increase over FR, suggesting the effect of each mutation in terms of **τ** was additive in the triple mutant. FR-MQV closely follows the spectral characteristics of FR-Q and FR-MQ, all of which have reduced spectral widths in absorbance and emission spectra with blue shifts in the absorption spectra with ~20% or higher peak absorbance with respect to FR. Consistent with the Strickler–Berg relationship, we calculate higher rate constants for radiative decay in FR mutants with the M42Q mutation. Based on these values, we successfully engineered the decrease of k_non-rad_ with a simultaneous increase of k_rad_ in FR, which is an efficient way of engineering the molecular brightness of an FP. FR-MQV has a k_rad_ comparable to the brightest RFP to date - mScarlet and its cellularly brighter mutant mScarlet-I. Finally, these mutations translate to high cellular brightness without compromising the physiological properties of the parent FR, as seen from the cytotoxicity, maturation and imaging assays performed in this study.

Crystal structures indicate that the phenol group of FP chromophores can occupy either the trans (non-fluorescing) or the cis (fluorescing) configuration^52,53^ and in FR, the latter can potentially form a hydrogen bond with the hydroxy group of the S144 residue (Figure 2).^51^ The location of C159 suggests that it may play a similar role for the trans form of the chromophore. The position 159 was recently discussed to be critical in terms of molecular brightness for FR.^51^ The propensity to switch to the trans form may be reduced when the residue at this position is substituted for an aliphatic group. The cis to trans isomerization is a reversible process and is manifested as the reversible component of the photobleaching trace under continuous and pulsed (ms to s) illumination.^45,46^ The reduction in dark state conversion in the photobleaching traces of FR-C159V and FR-C159L indicate that there is a reduction in the tendency of the chromophore to switch to a dark, trans form. FR-C159V also did not show a gradual photo-activation of fluorescence.^54^ This further suggests the chromophore might be locked into one configuration. Leu has a larger aliphatic sidechain than Val; thus, the shorter lifetime and complicated photo-activation behavior of FR-C159L may be a steric effect, with Val providing a better spatial fit that restricts chromophore movement into the dark, trans state.

Time-resolved ultrafast spectroscopy previously demonstrated that TagRFP-675 and mKate-Q display large Stokes shifts because there are multiple emissive species in the form of non-interconverting hydrogen bonded conformers.^37,38^ Multiple emissive species tend to broaden spectra and correlate with lower **ϕ** and shorter **τ** values.^38^ Steady state excitation-dependent emission spectra and fluorescence lifetime decay spectra collected in multiple emission windows dismiss the existence of multiple emissive species in M42Q mutants of FR (Details in Supplementary Information S4f: Figure S6 and S4g: Table S4). This is consistent with the minimal change in Stokes shift in the M42Q mutants of FR in comparison to mKate-Q. Gln has a polar sidechain accessible to different rotameric states.^55^ It is possible that this residue at position 42 rearranges itself into a conformation that might restrict the number of emissive species. This is supported by UV-Vis spectral data. which reveal that incorporating M42Q decreases the spectral width, blue shifts the absorption and emission spectra and increases the radiative rate constant. Other than Gln, we observed red fluorescence from the mutant with Ile at this position. Ile is of similar size to Gln and Met but is aliphatic in chemical nature. Characterization of the FR-MI mutant revealed a relative decrease in **ε** and **ϕ** with a decrease in **τ** with respect to the parent FR-M. We also observed an undesirable green emitting chromophore in the FR-MI and mKate-Q mutants, which was absent from the M42Q mutants of FR (Supplementary Information S4e: Figure S5), suggesting complete maturation to a red chromophore in FR-42Q mutants. Alkali denaturation of FR-Q, FR-MQ and FR-MQV shows cleavage of the chromophore only from the β-carbon of the Tyr sidechain, which is indicative of base access to a single site for hydrolysis (Supplementary Information S4d: Figure S4). The β-carbon of the Tyr sidechain in RFPs is spatially distant (>10Å based on crystal structure data) from the acylimine end of the chromophore. These molecular and photophysical properties suggest that factors other than steric bulk may be responsible for the changes observed in the M42Q mutants of FR.

The ~5-fold increase in brightness observed in FR-MQV relative to FR (Table 2) is likely an amalgamation of the higher molecular brightness of the M42Q and the C159V mutations along with the higher cellular brightness seen for FR-M. The trends for brightness measured through cytometry and microscopy are similar, but FACS measurements employ short laser exposure times with fast, sensitive detectors like photomultiplier tubes (PMTs). In assays involving microscopy, cells are subject to relatively longer exposure times and higher irradiances (~ms timescales) imposed by acquisition times of cameras, which can lead to photobleaching. The discrepancy between the absolute brightness values between FACS and confocal microscopy measurements were also observed for the control RFP mScarlet.

**Table 2:** Brightness measurements (from FACS) in HeLa and bacterial cells indicate that improvements in molecular and cellular brightness occur in tandem. On normalizing the cellular brightness with respect to FR-M, the brightness seen in FR-MQV seems to be due to an additive effect of the increase in molecular and cellular brightness.

| Protein | Molecular<br>Brightness<br>($\epsilon_{\max} * \phi$) | Bacterial Cell<br>Brightness<br>(585±15 nm) | HeLa Cell<br>Brightness<br>(585±15 nm) | Normalized relative to FR-M | |
| --- | --- | --- | --- | --- | --- |
|  |  |  |  | Molecular<br>Brightness | HeLa Cell<br>Brightness |
| **FR** | 1 | 1 | 1 | 0.9 | 0.5 |
| **FR-M** | 1.2 | 1.3 | 2.1 | 1 | 1 |
| **FR-Q** | 2 | 1.4 | - | 1.7 | - |
| **FR-MQ** | 2.6 | 1.7 | 3.6 | 2.3 | 1.7 |
| **FR-MQV** | 3.4 | 1.9 | 5.3 | 2.9 | 2.5 |

All three mutations in FR-MQV face into the protein β-barrel and therefore seem to minimally perturb the cellular properties of FR. The high scores for the OSER and Golgi-localization assays for all FR mutants indicate physiological properties of FPs are not appreciably affected by mutations which have side chains facing into the protein β-barrel. In the development of FR from mKate and mKate2, most of the mutations that optimized the FP for biological properties were external.^36^ We targeted positions to increase brightness, and thus we did not alter these external positions. Consequently, the mutants retained FR’s original performance in terms of localization and cytotoxicity. Though chromophore maturation kinetics can be greatly altered with internal mutations,^34^ in this case, the chromophore maturation was not substantially slowed down by these mutations. Figure 6 is a schematic indicating the postulated effect of each mutation in the development of FR-MQV.

**Figure 6:**
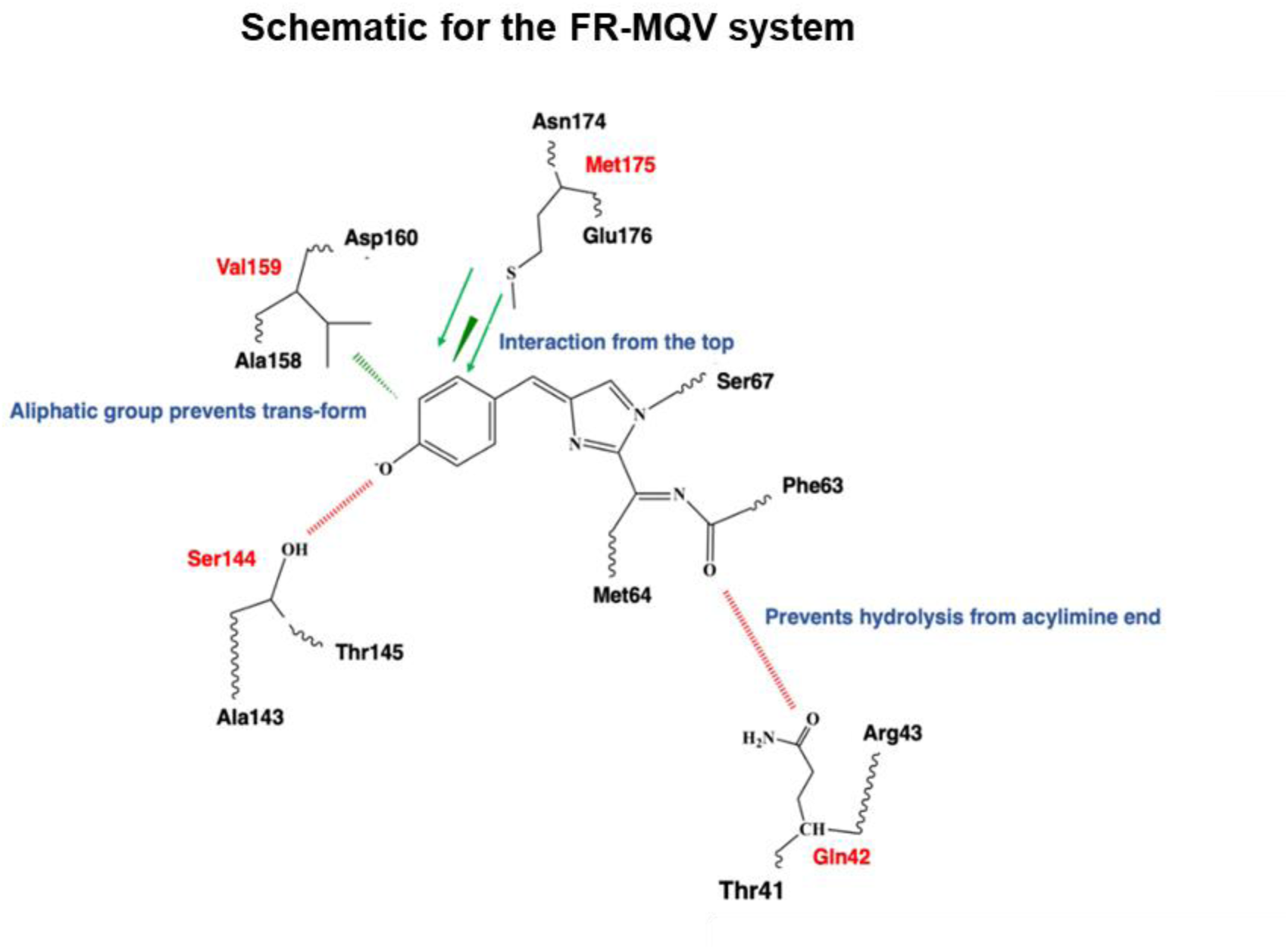
The overall molecular schematic describing the changes in FR-MQV. The L175M mutation from FR-M sits on top of the chromophore. The C159V mutation may reduce the propensity of the cis-trans isomerization of the chromophore, while the M42Q mutation possibly interacts with the acylimine end of the chromophore.

## Conclusions

In this study we present the development of FR-MQV, an FP with high brightness that maintains favorable cellular properties of FR. The C159V mutation appears to play a critical role in restricting the dark state conversion of the protein and increasing fluorescence lifetime, which leads to an increase in **ϕ**. The M42Q mutation plays an important role in enhancing the molecular brightness by increasing both the **ϕ** and **ε**. Interestingly, this mutation had different effects in closely related FPs including mKate and TagRFP-675. The previously reported L175M mutation contributes to robust expression levels which improves cellular brightness for FR mutants compared to FR. With its increased brightness and favorable cellular properties, FR-MQV shows promise as a template for further rounds of engineering suited to specific imaging applications like FLIM and FRET. Increasing fluorescence lifetime through further rounds of engineering may increase quantum efficiencies of emission to values near mScarlet with further reduction in non-radiative rates.

## Acknowledgements

SM was supported by the NIH/CU Molecular Biophysics Training Program (T32). This work was supported by the NSF Physics Frontier Center at JILA (PHY 1734006 to R.J.) and NIH DP1 GM114863 (to A.E.P). We thank Dr. Liya Muslinkina at the Macromolecular Crystallography Laboratory of the Frederick National Laboratory for Cancer Research, Argonne for helpful discussions about the FR crystal structure. We acknowledge Dr. Joe Dragavon, Theresa Nehreini, Dr. Evan Pratt, Dr. Maria Lo, Pia Friis and David Simpson for their invaluable help with this project. We also express our gratitude towards Shambojit Roy of the Cha-Goodwin Lab at the Chemical and Biological Engineering Department of CU Boulder for the kind gift of chloramphenicol. RJ is a staff member in the Quantum Physics Division of the National Institute of Standards and Technology (NIST). Certain commercial equipment, instruments, or materials are identified in this paper in order to specify the experimental procedure adequately. Such identification is not intended to imply recommendation or endorsement by the NIST, nor is it intended to imply that the materials or equipment identified are necessarily the best available for the purpose. We acknowledge the Flow Cytometry facility at BioFrontiers Institute, CU Boulder (grant # S10ODO21601). The imaging work was performed at the BioFrontiers Institute Advanced Light Microscopy Core facility. Spinning disc confocal microscopy was performed on Nikon Ti-E microscope supported by the BioFrontiers Institute and the Howard Hughes Medical Institute. Spectroscopy was carried out at the JILA shared facility in the W. Keck Lab.

## Methods and materials

No unexpected or unusually high safety hazards were encountered.

### Mutagenesis, cloning and construct development

#### Yeast Constructs

The QuickChange site-directed mutagenesis method was used for making point mutations using PfuTurboDNA polymerase and a thermocycler. Libraries with multiple site-directed targets were created using a splicing overlap extension reaction. Primers were designed to introduce the desired mutations and the initial PCRs generate overlapping gene segments that are used as template DNA for another PCR to create a full-length product. Fresh competent yeast cells (*Saccharomyces cerevisiae* BY4741) were prepared prior to electroporation. Cells, DNA and cut pYestDest52 vector were combined and left on ice for 5 min. Electroporation conditions (Bio-Rad Gene Pulser Xcell) were as follows: C = 25 μF, PC = 200 Ohm, V = 1.5 kV (in 0.2 cm cuvettes). Cells were passed twice prior to expression. Mutants were transferred to the pBad-His vector for expression/Ni-NTA protein purification.

#### Bacterial Expression Constructs

For bacterial expression, FR-Q and FR-MQ were made using the Q5 Site-Directed Mutagenesis Kit (New England Biolabs) with pBad-FR and pBad-FR-M, respectively, as the templates and the primers tCGCCACACAGGACACAAG and cCACATATGTCTCATCGTCAGC. FR-MQ C159V was also made via Q5 mutagenesis with the above primers and FR-M C159V as the template. The M42Q mutation was also made in pBad-mKate via Q5 mutagenesis and the corresponding A44Q mutation was made in pBad-mScarlet and pBad-mCherry using overlap extension PCR.

#### Mammalian Expression Constructs for FACS

FR-MQ and FR-MQV were expressed as histone H2B fusion proteins in HeLa cells. Specifically, the FR-MQ and FR-MQV mutants were PCR amplified from the pBad constructs with the upstream primer GGTATGGCTAGCATGACTGGTG and a reverse primer that introduces a NotI site adjacent to the stop codon (acatGCGGCCGCTCATTTCCCTCCATC). Similarly, mScarlet-I (mScarlet T74I) was PCR amplified from a Q5-generated pBad construct (primers AGGGCCTTCAtCAAGCACCCC and GCAGCCGTAC-ATGAACTGAGG) with the same upstream primer and a reverse primer that introduces an XcmI site adjacent to the stop codon. The FR-MQ and FR-MQV products were cut with BamHI and NotI and ligated into BamHI/NotI cut piggyBac-H2B and the mScarlet-I product was cut with BamHI and XcmI and ligated into BamHI/XcmI cut piggyBac-H2B.

#### Mammalian Expression Constructs for OSER

To assess protein aggregation in mammalian cells, the mutants were expressed as fusions with CytERM in U2OS cells. FPs were amplified from the pBad construct with an upstream primer that introduces an AgeI site upstream of the start codon (gcatACCGGTCGCCACCATGGTGTCCG-AGCTGATTAAGG) and the reverse primer described above that introduces a NotI site. The AgeI/NotI cut PCR product was ligated into AgeI/NotI cut CytERM vector and transfected into U2OS cells as described above for HeLa cells.

#### Mammalian Expression Constructs for GalT-FP fusions

To localize our mutants to the Golgi, they were expressed as GalT fusions in U2OS cells. The BamHI/NotI cut FR, FR-M and FR-MQV PCR products described above were ligated into BamHI/NotI cut GalT vector. mScarlet-I was cut from pBad with BamHI and SalI and ligated into BamHI/SalI cut GalT-mScarlet.

### Microfluidic based selection from site directed libraries

#### Library Targets

We targeted the positions 159, 161, 196 and 198 in FR (using FR numbering; Supplementary Information S1), expressed the libraries in yeast (*Sacchromyces cerevisiae*) and screened this library on a lifetime flow cytometer.^35^ Screening revealed the presence of brighter clones with longer and shorter lifetime than parent FR (lifetime ~ 2.05 ns). We performed another two rounds of FACS enrichment on a BD FACSAria Fusion Cell Sorter to remove the dim/non-fluorescent clones. Position 159 was mutated to I, L, V, F, M, C, A, G, T, S, W and R; position 161 was mutated to I, L, M, Q, N, H and K; position 196 was mutated to I, V, A and T; and position 198 was mutated to all possible amino acids. The library size was ~ 7,000 clones.

#### Post-Microfluidic Sorting

After selection, the collected yeast cells were grown in liquid culture and then plated, and 25 clones with unique lifetimes were picked from these sorter-enriched libraries using a lifetime assisted plate based screen discussed in a previous work.^35^ The FR site-directed (FSD) clones with unique DNA sequences were further characterized (Supplementary Information S2: Table S2).

### Cell growth, transformation/transfection and protein purification protocols

#### Bacterial Transformation and Growth

Bacteria (competent *E. coli*, Top10 strain) were transformed with the DNA encoding an FP of interest in the pBad-His vector. Roughly 2–5 μL of DNA (~80 ng/mL concentrations) were slowly pipetted into ~50 μL of competent cells (Invitrogen). The cells were left on ice for 20 minutes, heat shocked for 45 seconds at 42ºC and then grown in antibiotic -free medium for 45 minutes. The cells were then plated (~25**–**50 μL) onto ampicillin-containing LB agar plates and grown at 37ºC overnight. Colored colonies were picked from these plates and grown in 2XYT medium containing ampicillin overnight at 37ºC and 230 RPM. The next morning, 1 mL of this culture was added to 100 mL of fresh 2XYT with ampicillin, grown for 3 hours at 37ºC to achieve an OD of ~0.6, and then 1 mL of 20% arabinose was added to the culture to initiate expression. The temperature of incubation was lowered to 28ºC to slow down bacterial growth and help protein folding and chromophore maturation. Depending on the maturation rate of the FP, they were grown at this temperature for 20**–**30 hours.

#### Bacterial Cell Lysis and Protein Purification

The induced cell cultures were spun down at 8,000 RPM for 20 minutes at 4ºC and the cell pellet frozen at −30°C to ease lysis. B-PER Bacterial Protein Extraction Reagent (ThermoFisher) was used to lyse the cells in the presence of protease inhibitor. The cells were lysed for 1 hour at room temperature and then spun down at 11,000 RPM and 4ºC for 15 and then 20 minutes to remove the cellular debris. The supernatant containing the 6x-His-tagged protein was filtered with a 0.45 μm polyethersulfone membrane syringe filter and incubated with Ni-NTA agarose for 1 hour on ice. The resin was then loaded into a column, washed with 10 and 20 mM imidazole and then the proteins eluted with 250 mM imidazole. The imidazole was removed using PD-10 desalting columns (GE HealthCare) or 24 hours of dialysis using SnakeSkin dialysis tubing (ThermoFisher) into Tris-HCL buffer (pH 7.4) or saline PBS buffer. These samples were used for *in vitro* photophysical analyses.

#### Mammalian cell Growth and Transfection

HeLa/U2OS cells were cultured in RPMI medium (Gibco Life Technologies) supplemented with penicillin/streptomycin (Gibco Life Technologies) and 10% heat-inactivated fetal bovine serum (Sigma-Aldrich) at 37°C with 5% CO2 plus humidity. For imaging experiments, U2OS cells were grown in 35 mm imaging dishes (made in-house from Corning 35 x 10 mm dishes with VWR 18 x 18 mm #1.5 cover slips). All CyTERM constructs were transiently transfected for 18–24 hours using Lipofectamine 3000 (Invitrogen) or TransIT-LT1 transfection reagent (Mirus) according to the manufacturer’s instructions. For FACS experiments, we used HeLa cells transiently transfected using the TransIT-LT1 reagent (Mirus, catalog #MIR2304) and prepared for FACS analysis after 48 hours.

### *In vitro* photophysical measurements

#### Instrumentation

Absorption spectra were collected on a Cary5000 UV-Vis Near IR Spectrophotometer using a double beam mode with matched cuvettes and blank subtraction. Samples were diluted using 1X -Tris-HCl Buffer (pH ~7.4) and absorbance was measured at optical densities (ODs) between 0.05 and −0.25 to maintain measurements in the linear range of the Beer–Lambert’s law. Fluorescence measurements were performed with a HORIBA Jobin Yvon Fluorolog-3 FL3-222.

#### Extinction Coefficient (ε_max_) Measurements

Alkali denaturation was used to estimate the ratiometric values of **ε**_**max**_ for the samples. The main text of the article reports an average of three or more independent measurements performed for FPs reported with a standard deviation error. FPs with fewer measurements are reported without an error bar.

#### To measure the ε_max_, the following protocol was used

a)Blank1: 900 μL Tris-HCl buffer (pH 7.4) spectrum was recorded.; (b)Blank2: 900 μL Tris-HCl buffer (pH 7.4) +100 μL 10 M NaOH (pH~14); (c) 900 μL Tris-HCl buffer (pH 7.4) + a few μL of concentrated pure protein sample was added to adjust the absorbance to a value of OD ~0.1. A spectrum from 300–700 nm was recorded.; (d) 100 μL of 10 M NaOH was added to this solution and a spectrum in the same range was recorded immediately. Kinetic effects start playing a role on delaying the spectral measurement as degradation product peaks are known to drift in amplitude and wavelength over time.

#### Calculation of Extinction Coefficient (ε_max_)

We performed titration-based ε_max_ calculations for FR-M, FR-MQV and mScarlet-I. The numbers obtained compare well to the values measured by the one-step alkali denaturing method.16 The values were based on the mathematical relationship stated below, as FPs of the FR family are known to exhibit backbone cleavage.^36^ To verify if this method was valid for FR-MQV we also performed SDS-PAGE, where purified proteins (~10 μg) were run on TruPAGE precast 4–20% gradient acrylamide gels (Sigma-Aldrich) in TEA–Tricine running buffer with the Spectra Multicolor Broad Range protein ladder (ThermoFisher). (Supplementary Information S6a: Figure S12) For other FPs that do not exhibit backbone cleavage, like mScarlet, the absorption at 380 nm was disregarded.

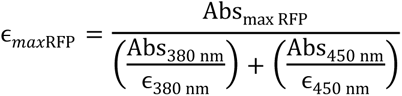

#### Quantum Yield Measurements

Freshly prepared or flash frozen purified protein was diluted with Tris-HCl buffer (pH 7.4) in a 1 cm path length Quartz cuvette to an OD of ~0.1. A matched cuvette was used for baseline correction to measure absorption spectra. The same cuvette with the solution in it was transferred to the fluorimeter for collecting fluorescence spectra. After each absorption and emission scan, 200–250 μL of the sample was removed and replaced with fresh buffer to create a step dilution. This step dilution was repeated 4–5 times for each sample. RFPs were excited at 520 nm such that the entire emission spectrum was recorded for each FP (even for blue-shifted RFPs) with high enough absorption. For each FP, integrated fluorescence was calculated for the area under the RFP emission feature on the emission spectra and was plotted against the corresponding OD at 520 nm from absorption spectrum. The integrated fluorescence versus OD plot can be fitted with a straight line (See Supplementary Information S6b: Figure S13) of the form: y = *slope × x*, where *y* is the integrated fluorescence and *x* is the OD. ϕ of the sample is computed as: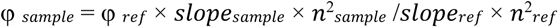 Where the refractive index of the sample and the reference are *n*_sample_ and *n*_ref_, respectively. mCherry (ϕ = 0.22 in Tris-HCl, pH 7.4), Cresyl Violet (ϕ = 0.54 in EtOH) and mScarlet (ϕ = 0.72 with respect to mCherry) were used as references.^16^ Multiple measurements were taken for each sample. (Supplementary Information S6b: Table S9)

#### pKa Measurements

pKa measurements were performed by preparing buffers in the range of pH 2 to 12. The pH was measured for each buffer to confirm the calculated pH values using a pH meter. Fixed amounts of concentrated pure protein samples were added to 1 mL buffer in a quartz cuvette with a 1 cm path length and a fluorescence spectrum was recorded in each case with excitation at 520 nm. The maximum value of fluorescence counts was used to normalize the fluorescence spectra for each protein. The data were plotted and was fit to the sigmoidal curve shown in Supplementary Information S4h: Figure S7. The calculated pKa values and reported values for published RFPs have been provided in Supplementary Information S4h: Table S5. Buffers in the pH range 2–3 were prepared with a dilution of glycine and 1 M HCl; in the range 3–6 were prepared using dilutions of 0.1 M citric acid and 0.1 M Na-citrate; in the range 6–8 were prepared with dilutions of 0.2 M Na2HPO4 and 0.2 M KH2PO4; in the range of 9–12 were prepared with dilutions of 0.2 M glycine and 1 M NaOH and Tris-HCl buffer was used for the pH 7.4 measurement.

#### Lifetime Measurements Based on the Lifetime Flow Cytometer

The lifetime flow cytometer utilizes frequency-domain phase fluorimetry to select analytes based on excited state lifetime. The details of this set-up are discussed in our previous publications.^35,56^ The excitation beam is sinusoidally modulated at 29.5 MHz. The fluorescence trace closely follows the excitation trace in frequency but with a lower modulation depth and a phase shift that corresponds to the average time spent by the analyte in the excited state (average lifetime). A high-speed lock-in amplifier determines the phase shift, which is then converted to a lifetime value in the time domain. Lifetime_analyte_ = Lifetime_reference_(ns) + tangent (Phase-Shift) /29.5 MHz For the reference, mCherry (~1.6 ns) was used. FR had a mean lifetime 2.05 ± 0.15 ns on this device.

#### Steady-State Lifetime Measurements for Pure Proteins and Lysates

All lifetime measurements on purified proteins or filtered cell lysates were performed on a commercial TCSPC system (Fluoro-time 100, PicoQuant) using a 560 nm pulsed laser diode head excitation source with a repetition rate of 5 MHz. Emission was collected either using a red filter set (600 ± 30 nm) or a far-red filter set (670 ± 30 nm), to check for multiple species in the excited state or interconverting forms. The instrument response function (IRF) was collected using a Ludox (Millipore Sigma) colloidal silica, whereas the samples were diluted to an OD value <0.05 for the measurement. A minimum of 20,000 photon counts were used to generate the fluorescence decays. The fluorescence transient decays were fit to an iterative re-convolution with a bi-exponential function (or-tri exponential depending on the protein). The amplitudes and the components of the fits are provided in Supplementary Information S4g: Table S4.

#### Photobleaching Measurements

Photobleaching measurements were performed using an LED excitation source (Lumencor) on an Olympus IX-73 fluorescence microscope with *E. coli* cells expressing the FP of interest. Bacteria on plates were washed and dispersed in aqueous blank buffer containing 0.17% (w/v) yeast nitrogen base (Sigma-Aldrich) and 0.5% (w/v) ammonium sulfate (Sigma-Aldrich) then photobleached with excitation rate-normalized LED light. To bleach the sample, a 560 nm LED was used, whereas for repopulation with blue light, a 438 nm LED light source was used. Details of the measurements are discussed in Supplementary Information S4j: Figure S9.

#### Brightness in E. coli

To assess the cellular performance of the FR clones in this study, we performed a bacterial brightness assay at the single cell level on a droplet microfluidic sorting platform.^56^ Two biological replicates with three independent technical triplicates were performed for each measurement. The bacteria were grown, and expression was induced as previously described, cytometry was carried out at ~20–22 hours after starting induction. Each technical replicate involved 10,000 cells. Details of the screening protocol have been discussed in a previous publication.^56^

### Cellular brightness assays

#### Brightness in HeLa Cells

*(a) Flow Cytometry*—The proteins of interest in the FR family, along with some standard RFPs including mScarlet, mScarlet-I and mCherry, were fused to histone H2B and expressed in HeLa cells. Single-cell brightness was assessed by selecting single healthy cells based on forward and side-scattering photon counts on a BD FACSCelesta single cell analyzer after 48 hours of transfection. Untransfected cells were used as a control to background subtract and analyze the fluorescence in the red and green channels for the proteins of interest. In most cases (except mScarlet-I with only one biological replicate) three or more biological replicates with three technical replicates of each were analyzed to determine the mean fluorescence with standard deviation based on the number of measurements (Figure 4a and Table 2). The samples were excited by a 561 nm laser line for collecting red fluorescence through the TRITC filter set (585/30 nm) and a 488 nm laser line for collecting through a GFP filter set (530/30 nm). The residual fluorescence from the green channel was effectively at the background level of the EGFP-H2B control (displaying signal values ~20-fold higher than mScarlet with the highest green fluorescence in the series). Brighter red mutants had higher green fluorescence background, suggesting red fluorescence bleeding through the green channel. (b) *Confocal Microscopy—*HeLa cells grown in 35 mm imaging dishes (made in-house from Corning 35 x 10 mm dishes with VWR 18 x 18 mm #1.5 cover slips) were imaged 48 hours post transfection to maintain consistency with the FACS measurements. Before imaging, cells were washed three times with 2 mL phosphate-free HEPES-buffered Hanks’ balanced salt solution (HHBSS) containing 20 mM HEPES (Sigma), pH 7.4 and resuspended in 1.5 mL of the same buffer. Imaging was performed on a Nikon Ti-E spinning disc confocal microscope system. The imaging dishes were mounted on the microscope in an environmentally controlled chamber (Oko Labs; set to 37°C, 5% CO_2_, 90% humidity) and viewed with a 40x (NA 0.95) air objective. A 560 nm laser was used for illumination and a 590–650 nm band pass filter (TRITC) was used for the detection of fluorescence with 200 ms exposure time and 20% laser power of the instrument. To minimize photobleaching, the focus of the microscope was adjusted to a lower laser power (5%) at only the center spot of the large image. Prolonged exposure of FR and its mutants to laser light can lead to lower brightness, which was a critical factor for this assay. Large images (~1600 x 1000 *μm*) were taken for each dish for each replicate.

#### Data Analysis for Brightness Assays Using Confocal Microscopy

The imaged cells were analyzed using the suite CellProfiler.^57^ A pipeline was created that would identify objects that are above the noise background. A binarized image was thus created, then a gate to filter objects typical of the size of nuclei was selected (2–25 *μm*), which selected the fluorescing nuclei in the H2B construct. The filtered objects in the parent cells were then quantified for mean brightness. Untransfected cells with cellular auto-fluorescence were the first to be analyzed, which gave us the estimation of bleed-through fluorescence.

### Cellular localization assays

#### Imaging Conditions

U2OS cells were transfected with the constructs of interest as described previously. The cells were imaged 24 hours post transfection with a Nikon Ti-E spinning disc confocal microscope. Several large images (~1,680 × 1,000 *μm*) were captured while scanning the z-focus (z-stacks) for optimum focusing of all the cells appearing in the field of view. Typical scanned depth was 5.0–7.8 *μm* and was evenly separated into 5–7 layers. Maximum intensity projection of the z-stacks was used for the quantification of OSER score. For GalT scoring, Z-stacks and independent frames were used for quantification of localization of FPs to the Golgi.

#### Data Analysis

The data analysis for the quantification of OSER scores was done in accordance with our previous report.^35^ In brief, the suite CellProfiler was used to develop a pipeline that can identify cells based on size (20–50 μm), shape and fluorescence intensity. The program identifies oligomerization sub-structures in these cells, filters them and then quantifies them as whorls. The pipeline further continues to relate the number of whorl structures with each cell and a MATLAB code quantifies the number of perfect and imperfect (cells with whorls). A score of 100 indicates 100% of the cells are whorl free and 0% indicates all cells have whorls in them. FR and its mutants display a high OSER score, whereas TagRFP-T, was used as a negative control. The imaging data for the GalT-FP construct was analyzed using a blinding approach where multiple individuals were provided with image sets that were randomized. Individual images were scored as having fluorescence signal localized to the Golgi (characterized by a fist-like structure near the nucleus) or not (Figure 5b). The approximate size of a healthy nucleus was estimated using ImageJ.^58^ If the cells expressed FP outside this fist like structure in the form of puncta or just smeared across the cytoplasm, they were considered mis-localized.

### Chromophore maturation kinetics and cytotoxicity

#### Cytotoxicity Assay

HeLa cells were transfected with the H2B-FP fusion constructs as discussed previously. Two biological replicates were prepared for each FP. Day 2 was defined as 48 hours post transfection. RFP and EGFP transfected cells were mixed at a 50:50 ratio by volume. A sample of the cell mixture was prepared for a flow cytometry measurement (BD FACSCelesta) to determine the actual RFP:EGFP ratio. The rest of the mixture was re-plated for further growth and proliferation. Independent screens of just EGFP cells and RFP cells were carried out first. Cells with green fluorescence higher than background were classified as “EGFP” and cells with red fluorescence higher than background were classified as “RFP”. On Day 6, the re-plated cells were again subjected to flow cytometry to quantify the change in the RFP and EGFP populations and determine a relative level of cytotoxicity. The cytotoxicity score was then calculated as the change in the RFP:GFP ratio for each sample. EGFP was used for normalization because it has been shown to be minimally cytotoxic. 59 The data for the assay are presented in Supplementary Information S5b: Figure S10.

#### Maturation Kinetics

Bacteria (Top 10, *E. coli*) expressing FPs were grown in expression media for 4 hours (28ºC, 230 RPM) to facilitate FP production in the exponential growth log phase. Chloramphenicol (250 μg/mL) was added to the cultures to stall bacterial growth and protein production.^60^ The cultures were maintained at 37ºC and 230 RPM and aliquots were taken every 30-60 minutes to measure the Optical Density (OD) and fluorescence. The t1/2 at 37ºC was calculated by calculating the time required for the FP to reach half the maximum fluorescence value. The data for the assay is presented in Supplementary Information S5c: Figure S11 and Table S8.

