## Supplementary Information for "Structure-guided point mutations on FusionRed produce a brighter red fluorescent protein"


| Section | Title |
|---|---|
| S1 | Amino acid sequence information |
| S2 | Amino acid sequence results from the site-directed library |
| S3 | Additional structural information |
| S4 | Additional photophysical and biochemical characterization |
| S5 | Cellular Assays: brightness, maturation and cytotoxicity |
| S6 | Extinction coefficient and quantum yield estimation |
| S7 | FR evolution table |

**S1: Amino acid sequence information:** Sequence information for the FR family with respect to the parental clones from the eqFP578[1] family. The amino acid sequences of avGFP[2] and DsRed are also listed in the table for comparison. The relevant mutations in this study have been highlighted in gold. The chromophore residues are shown in blue, internal mutations highlighted in grey and external mutations in white. All mutation position numbers in the text are with respect to FR numbering in this table.

| GFP Allignment | | | | | | | | | | 10 | | | | | | | | | | 20 | | | | | | | | | | 30 | | |
|---|---|---|---|---|---|---|---|---|---|---|---|---|---|---|---|---|---|---|---|---|---|---|---|---|---|---|---|---|---|---|---|---|
| avGFP | M | S | K | G | E | E | L | F | T | G | V | V | P | I | L | V | E | L | D | G | D | V | N | G | H | K | F | S | V | S | G | E |
| DsRed | M | R | S | S | K | N | V | I | K | E | F | M | R | F | K | V | R | M | E | G | T | V | N | G | H | E | F | E | I | E | G | E |
| eqFP578 | | | | M | S | E | L | I | K | E | N | M | H | M | K | L | Y | M | E | G | T | V | N | N | H | H | F | K | C | T | S | E |
| Katushka | | | | M | S | V | L | I | T | E | N | M | H | M | K | L | Y | M | E | G | T | V | N | D | H | H | F | K | C | T | S | E |
| TagRFP | | | | M | S | E | L | I | K | E | N | M | H | M | K | L | Y | M | E | G | T | V | N | N | H | H | F | K | C | T | S | E |
| mKate | | | | M | S | E | L | I | K | E | N | M | H | M | K | L | Y | M | E | G | T | V | N | N | H | H | F | K | C | T | S | E |
| mKate2 | | | M | V | S | E | L | I | K | E | N | M | H | M | K | L | Y | M | E | G | T | V | N | N | H | H | F | K | C | T | S | E |
| mKate2.5 | | | M | V | S | E | L | I | K | E | N | M | H | M | K | L | Y | M | E | G | T | V | N | N | H | H | F | K | C | T | S | E |
| FusionRed | | | M | V | S | E | L | I | K | E | N | M | P | M | K | L | Y | M | E | G | T | V | N | N | H | H | F | K | C | T | S | E |
| FusionRed-Q | | | M | V | S | E | L | I | K | E | N | M | P | M | K | L | Y | M | E | G | T | V | N | N | H | H | F | K | C | T | S | E |
| FusionRed-M | | | M | V | S | E | L | I | K | E | N | M | P | M | K | L | Y | M | E | G | T | V | N | N | H | H | F | K | C | T | S | E |
| FusionRed-MQ | | | M | V | S | E | L | I | K | E | N | M | P | M | K | L | Y | M | E | G | T | V | N | N | H | H | F | K | C | T | S | E |
| FusionRed-MQV | | | M | V | S | E | L | I | K | E | N | M | P | M | K | L | Y | M | E | G | T | V | N | N | H | H | F | K | C | T | S | E |
| FusionRed Alignment | | | 1 | | | | | | | | | 10 | | | | | | | | | | 20 | | | | | | | | | | 30 |

| | | | | | | | 40 | | | | | | | | | | 50 | | | | | | | | | | 60 | | | | | | | | | | 70 | | |
|---|---|---|---|---|---|---|---|---|---|---|---|---|---|---|---|---|---|---|---|---|---|---|---|---|---|---|---|---|---|---|---|---|---|---|---|---|---|---|---|
| G | E | G | D | A | T | Y | G | K | L | T | L | K | F | I | C | T | T | G | - | K | L | P | V | P | W | P | T | L | V | T | T | F | S | Y | G | V | Q | C | F |
| G | E | G | R | P | Y | E | G | H | N | T | V | K | L | K | V | T | K | G | G | P | L | P | F | A | W | D | I | L | S | P | Q | F | Q | Y | G | S | K | V | Y |
| G | E | R | K | P | Y | E | G | T | Q | T | M | K | I | K | V | V | E | G | G | P | L | P | F | A | F | D | I | L | A | T | S | F | M | Y | G | S | K | T | F |
| G | E | G | K | P | Y | E | G | T | Q | T | M | K | I | K | V | V | E | G | G | P | L | P | F | A | F | D | I | L | A | T | S | F | M | Y | G | S | K | T | F |
| G | E | G | K | P | Y | E | G | T | Q | T | M | R | I | K | V | V | E | G | G | P | L | P | F | A | F | D | I | L | A | T | S | F | M | Y | G | S | R | T | F |
| G | E | G | K | P | Y | E | G | T | Q | T | M | R | I | K | V | V | E | G | G | P | L | P | F | A | F | D | I | L | A | T | S | F | M | Y | G | S | K | T | F |
| G | E | G | K | P | Y | E | G | T | Q | T | M | R | I | K | A | V | E | G | G | P | L | P | F | A | F | D | I | L | A | T | S | F | M | Y | G | S | K | T | F |
| G | E | G | K | P | Y | E | G | T | Q | T | M | R | I | K | V | V | E | G | G | P | L | P | F | A | F | D | I | L | A | T | S | F | M | Y | G | S | K | T | F |
| G | E | G | K | P | Y | E | G | T | Q | T | **M** | R | I | K | V | V | E | G | G | P | L | P | F | A | F | D | I | L | A | T | S | F | M | Y | G | S | R | T | F |
| G | E | G | K | P | Y | E | G | T | Q | T | **Q** | R | I | K | V | V | E | G | G | P | L | P | F | A | F | D | I | L | A | T | S | F | M | Y | G | S | R | T | F |
| G | E | G | K | P | Y | E | G | T | Q | T | **M** | R | I | K | V | V | E | G | G | P | L | P | F | A | F | D | I | L | A | T | S | F | M | Y | G | S | R | T | F |
| G | E | G | K | P | Y | E | G | T | Q | T | **Q** | R | I | K | V | V | E | G | G | P | L | P | F | A | F | D | I | L | A | T | S | F | M | Y | G | S | R | T | F |
| G | E | G | K | P | Y | E | G | T | Q | T | **Q** | R | I | K | V | V | E | G | G | P | L | P | F | A | F | D | I | L | A | T | S | F | M | Y | G | S | R | T | F |
| | | | | | | | | | 40 | | 42 | | | | | | | | 50 | | | | | | | | | | 60 | | | | | | | | | | 70 |

| | | | | | | | 80 | | | | | | | | | | 90 | | | | | | | | | | 100 | | | | | | | | | | 110 | | | | |
|---|---|---|---|---|---|---|---|---|---|---|---|---|---|---|---|---|---|---|---|---|---|---|---|---|---|---|---|---|---|---|---|---|---|---|---|---|---|---|---|---|---|
| S | R | Y | P | D | H | M | K | Q | H | D | F | F | K | S | A | M | P | E | G | Y | V | Q | E | R | T | I | F | F | K | D | D | G | N | Y | K | T | R | A | E | V | K |
| V | K | H | P | A | D | I | P | - | - | D | Y | K | K | L | S | F | P | E | G | F | K | W | E | R | V | M | N | F | E | D | G | G | V | V | T | V | T | Q | D | S | S |
| I | N | H | T | Q | G | I | P | - | - | D | L | F | K | Q | S | F | P | E | G | F | T | W | E | R | I | T | T | Y | E | D | G | G | V | L | T | A | T | Q | D | T | S |
| I | N | H | T | Q | G | I | P | - | - | D | F | F | K | Q | S | F | P | E | G | F | T | W | E | R | I | T | T | Y | E | D | G | G | V | L | T | A | T | Q | D | T | S |
| I | N | H | T | Q | G | I | P | - | - | D | F | F | K | Q | S | F | P | E | G | F | T | W | E | R | V | T | T | Y | E | D | G | G | V | L | T | A | T | Q | D | T | S |
| I | N | H | T | Q | G | I | P | - | - | D | F | F | K | Q | S | F | P | E | G | F | T | W | E | R | V | T | T | Y | E | D | G | G | V | L | T | A | T | Q | D | T | S |
| I | N | H | T | Q | G | I | P | - | - | D | F | F | K | Q | S | F | P | E | G | F | T | W | E | R | V | T | T | Y | E | D | G | G | V | L | T | A | T | Q | D | T | S |
| I | N | H | P | Q | G | I | P | - | - | D | F | F | K | Q | S | F | P | E | G | F | T | W | E | R | V | T | T | Y | E | D | G | G | V | L | T | A | T | Q | D | T | S |
| I | K | H | P | P | G | I | P | - | - | D | F | F | K | Q | S | F | P | E | G | F | T | W | E | R | V | T | T | Y | E | D | G | G | V | L | T | A | T | Q | D | T | S |
| I | K | H | P | P | G | I | P | - | - | D | F | F | K | Q | S | F | P | E | G | F | T | W | E | R | V | T | T | Y | E | D | G | G | V | L | T | A | T | Q | D | T | S |
| I | K | H | P | P | G | I | P | - | - | D | F | F | K | Q | S | F | P | E | G | F | T | W | E | R | V | T | T | Y | E | D | G | G | V | L | T | A | T | Q | D | T | S |
| I | K | H | P | P | G | I | P | - | - | D | F | F | K | Q | S | F | P | E | G | F | T | W | E | R | V | T | T | Y | E | D | G | G | V | L | T | A | T | Q | D | T | S |
| I | K | H | P | P | G | I | P | - | - | D | F | F | K | Q | S | F | P | E | G | F | T | W | E | R | V | T | T | Y | E | D | G | G | V | L | T | A | T | Q | D | T | S |
| | | | | | | | | | | | 80 | | | | | | | | | | 90 | | | | | | | | | | 100 | | | | | | | | | | 110 |

| | | | | | | | | | | | | | | | | | | | | | | | | | | | | | | | | | | | | | | | | |
|---|---|---|---|---|---|---|---|---|---|---|---|---|---|---|---|---|---|---|---|---|---|---|---|---|---|---|---|---|---|---|---|---|---|---|---|---|---|---|---|---|
| | | | | | 120 | | | | | | | | | | 130 | | | | | | | | | | 140 | | | | | | | | | | 150 | | | | | |
| F | E | G | D | T | L | V | N | R | I | E | L | K | G | I | D | F | K | E | D | G | N | I | L | G | H | K | L | E | Y | N | Y | N | S | H | N | V | Y | I | M | A |
| L | Q | D | G | C | F | I | Y | K | V | K | F | I | G | V | N | F | P | S | D | G | P | V | M | Q | K | K | T | M | - | G | W | E | A | S | T | E | R | L | Y | P |
| L | Q | N | G | C | I | I | Y | N | V | K | I | N | G | V | N | F | P | S | N | G | S | V | M | Q | K | K | T | L | - | G | W | E | A | N | T | E | M | L | Y | P |
| L | Q | N | G | C | L | I | Y | N | V | K | I | N | G | V | N | F | P | S | N | G | P | V | M | Q | K | K | T | L | - | G | W | E | A | S | T | E | M | L | Y | P |
| L | Q | D | G | C | L | I | Y | N | V | K | I | R | G | V | N | F | P | S | N | G | P | V | M | Q | K | K | T | L | - | G | W | E | A | N | T | E | M | L | Y | P |
| L | Q | D | G | C | L | I | Y | N | V | K | I | R | G | V | N | F | P | S | N | G | P | V | M | Q | K | K | T | L | - | G | W | E | A | S | T | E | M | L | Y | P |
| L | Q | D | G | C | L | I | Y | N | V | K | I | R | G | V | N | F | P | S | N | G | P | V | M | Q | K | K | T | L | - | G | W | E | A | S | T | E | T | L | Y | P |
| L | Q | D | G | C | L | I | Y | N | V | K | I | R | G | V | N | F | P | A | N | G | P | V | M | Q | K | K | T | L | - | G | W | E | A | S | T | E | T | L | Y | P |
| L | Q | D | G | C | L | I | Y | N | V | K | V | R | G | V | N | F | P | A | N | G | P | V | M | Q | K | K | T | L | - | G | W | E | A | S | T | E | T | M | Y | P |
| L | Q | D | G | C | L | I | Y | N | V | K | V | R | G | V | N | F | P | A | N | G | P | V | M | Q | K | K | T | L | - | G | W | E | A | S | T | E | T | M | Y | P |
| L | Q | D | G | C | L | I | Y | N | V | K | V | R | G | V | N | F | P | A | N | G | P | V | M | Q | K | K | T | L | - | G | W | E | A | S | T | E | T | M | Y | P |
| L | Q | D | G | C | L | I | Y | N | V | K | V | R | G | V | N | F | P | A | N | G | P | V | M | Q | K | K | T | L | - | G | W | E | A | S | T | E | T | M | Y | P |
| L | Q | D | G | C | L | I | Y | N | V | K | V | R | G | V | N | F | P | A | N | G | P | V | M | Q | K | K | T | L | - | G | W | E | A | S | T | E | T | M | Y | P |
| | | | | | | | | | 120 | | | | | | | | | | 130 | | | | | | | | | | | 140 | | | | | | | | | | 150 |

| | | | | | | | | | | | | | | | | | | | | | | | | | | | | | | | | | | | | | | | | | |
|---|---|---|---|---|---|---|---|---|---|---|---|---|---|---|---|---|---|---|---|---|---|---|---|---|---|---|---|---|---|---|---|---|---|---|---|---|---|---|---|---|---|
| | | | | 160 | | | | | | | | | | 170 | | | | | | | | | | 180 | | | | | | | | | | 190 | | | | | | | |
| D | K | Q | K | N | G | I | K | V | N | F | K | I | R | H | N | I | E | D | G | S | V | Q | L | A | D | H | Y | Q | Q | N | T | P | I | G | D | - | G | P | V | L | L |
| R | - | - | D | G | V | L | K | G | E | I | H | K | A | L | K | L | K | D | G | G | H | Y | L | V | E | F | K | S | I | Y | M | A | K | K | P | - | - | - | V | Q | L |
| A | - | - | D | G | G | L | R | G | H | S | Q | M | A | L | K | L | V | G | G | G | Y | L | H | C | S | F | K | T | T | Y | R | S | K | K | P | A | K | N | L | K | M |
| A | - | - | D | S | G | L | R | G | H | S | Q | M | A | L | K | L | V | G | G | G | Y | L | H | C | S | L | K | T | T | Y | R | S | K | K | P | A | K | N | L | K | M |
| A | - | - | D | G | G | L | E | G | R | S | D | M | A | L | K | L | V | G | G | G | H | L | I | C | N | F | K | T | T | Y | R | S | K | K | P | A | K | N | L | K | M |
| A | - | - | D | G | G | L | E | G | R | S | D | M | A | L | K | L | V | G | G | G | H | L | I | C | N | L | K | T | T | Y | R | S | K | K | P | A | K | N | L | K | M |
| A | - | - | D | G | G | L | E | G | R | A | D | M | A | L | K | L | V | G | G | G | H | L | I | C | N | L | K | T | T | Y | R | S | K | K | P | A | K | N | L | K | M |
| A | - | - | D | G | G | L | E | G | A | C | D | M | A | L | K | L | V | G | G | G | H | L | I | C | N | L | E | T | T | Y | R | S | K | K | P | A | K | N | L | K | M |
| A | - | - | D | G | G | L | E | G | A | C | D | M | A | L | K | L | V | G | G | G | H | L | I | C | N | L | E | T | T | Y | R | S | K | K | P | A | T | N | L | K | M |
| A | - | - | D | G | G | L | E | G | A | C | D | M | A | L | K | L | V | G | G | G | H | L | I | C | N | L | E | T | T | Y | R | S | K | K | P | A | T | N | L | K | M |
| A | - | - | D | G | G | L | E | G | A | C | D | M | A | L | K | L | V | G | G | G | H | L | I | C | N | M | E | T | T | Y | R | S | K | K | P | A | T | N | L | K | M |
| A | - | - | D | G | G | L | E | G | A | C | D | M | A | L | K | L | V | G | G | G | H | L | I | C | N | M | E | T | T | Y | R | S | K | K | P | A | T | N | L | K | M |
| A | - | - | D | G | G | L | E | G | A | V | D | M | A | L | K | L | V | G | G | G | H | L | I | C | N | M | E | T | T | Y | R | S | K | K | P | A | T | N | L | K | M |
| | | | | | | | | | | 159 | 160 | | | | | | | | | | 170 | | | | | 175 | | | | | 180 | | | | | | | | | | 190 |

| | | | | | | | | | | | | | | | | | | | | | | | | | | | | | | | | | | | | | | | | | | |
|---|---|---|---|---|---|---|---|---|---|---|---|---|---|---|---|---|---|---|---|---|---|---|---|---|---|---|---|---|---|---|---|---|---|---|---|---|---|---|---|---|---|---|
| | | 200 | | | | | | | | | | 210 | | | | | | | | | | 220 | | | | | | | | | | 230 | | | | | | | | | | 240 |
| P | D | N | H | Y | L | S | T | Q | S | A | L | S | K | D | P | N | E | K | R | D | H | M | V | L | L | E | F | V | T | A | A | G | I | T | H | G | M | D | E | L | Y | K |
| P | G | Y | Y | Y | V | D | S | K | L | D | I | T | S | H | - | N | E | D | Y | T | I | V | E | Q | Y | E | R | T | E | G | R | H | H | L | F | L | | | | | | |
| P | G | F | H | F | V | D | H | R | L | E | R | I | K | E | - | A | D | K | E | T | Y | V | E | Q | H | E | M | A | V | A | K | Y | C | D | L | P | S | K | L | G | H | R |
| P | G | F | Y | F | V | D | R | R | L | E | R | I | K | E | - | A | D | K | E | T | Y | V | E | Q | H | E | M | A | V | A | R | Y | C | D | L | P | S | K | L | G | H | S |
| P | G | V | Y | Y | V | D | H | R | L | E | R | I | K | E | - | A | D | K | E | T | Y | V | E | Q | H | E | V | A | V | A | R | Y | C | D | L | P | S | K | L | G | H | K |
| P | G | V | Y | Y | V | D | R | R | L | E | R | I | K | E | - | A | D | K | E | T | Y | V | E | Q | H | E | V | A | V | A | R | Y | C | D | L | P | S | K | L | G | H | K |
| P | G | V | Y | Y | V | D | R | R | L | E | R | I | K | E | - | A | D | K | E | T | Y | V | E | Q | H | E | V | A | V | A | R | Y | C | D | L | P | S | K | L | G | H | R |
| P | G | V | Y | N | V | D | R | R | L | E | R | I | K | E | - | A | D | N | E | T | Y | V | E | Q | H | E | V | A | V | A | R | Y | S | T | G | G | A | G | D | G | G | K |
| P | G | V | Y | N | V | D | H | R | L | E | R | I | K | E | - | A | D | D | E | T | Y | V | E | Q | H | E | V | A | V | A | R | Y | S | T | G | G | A | G | D | G | G | K |
| P | G | V | Y | N | V | D | H | R | L | E | R | I | K | E | - | A | D | D | E | T | Y | V | E | Q | H | E | V | A | V | A | R | Y | S | T | G | G | A | G | D | G | G | K |
| P | G | V | Y | N | V | D | H | R | L | E | R | I | K | E | - | A | D | D | E | T | Y | V | E | Q | H | E | V | A | V | A | R | Y | S | T | G | G | A | G | D | G | G | K |
| P | G | V | Y | N | V | D | H | R | L | E | R | I | K | E | - | A | D | D | E | T | Y | V | E | Q | H | E | V | A | V | A | R | Y | S | T | G | G | A | G | D | G | G | K |
| P | G | V | Y | N | V | D | H | R | L | E | R | I | K | E | - | A | D | D | E | T | Y | V | E | Q | H | E | V | A | V | A | R | Y | S | T | G | G | A | G | D | G | G | K |
| | | | | | | | | | 200 | | | | | | | | | | 210 | | | | | | | | | | 220 | | | | | | | | | | 230 | | | 233 |

**Table S1:** Sequence data for the FR with respect to its parents (mKate and eqFP578), DsRed and avGFP. All mutations in the main text and the Supplementary Information have been described with respect to this table.

### S2. Amino acid sequence results from the site-directed library

***Selection of the C159V clone:*** The eight distinct mutants were screened for brightness in yeast cells in the green and red channel on a BD-FACS Celesta flow cytometer at the BioFrontiers Flow Cytometry core facility at the University of Colorado, Boulder. The green channel is an indicator for the undesirable immature green fluorescence peak seen in RFPs such as SDC-5 (an unpublished mCherry mutant with mutations of W143L, I161T, Q163C, I197R using mCherry sequence numbering). Filtered cell lysates of these mutants were also used for measuring lifetime on a time-correlated single photon counting (TCSPC) system that provided us with lifetimes to select the clones of interest. Sequence, lifetime and brightness data are summarized in Table S2.

**Table S2:** Sequence and screening results of the FR site-directed (FSD) library clones. The lifetime was measured using cell lysates on a TCSPC system. All fluorescence measurements were normalized to FR.

| Clones | 159 | 161 | Lifetime (ns) | Green fluorescence | Red fluorescence |
|---|---|---|---|---|---|
| FR | C | M | 1.78 | 100 | 100 |
| mCherry | - | - | 1.67 | 103 | 191 |
| SDC-5 | - | - | - | 811 | 421 |
| FR-1 | - | - | - | 185 | 352 |
| FSD-4 | T | M | 1.70 | 112 | 201 |
| FSD-5 | ? | ? | 1.94 | 100 | 193 |
| FSD-9 | V | M | 2.03 | 163 | 232 |
| FSD-11 | L | M | 1.21 | 107 | 177 |
| FSD-14 | A | H | 1.01 | 147 | 68 |
| FSD-15 | T | M | 1.73 | 105 | 134 |
| FSD-18 | T | I | 1.70 | 111 | 154 |
| FSD-19 | G | M | 1.94 | 105 | 87 |

### S3: Additional structural information

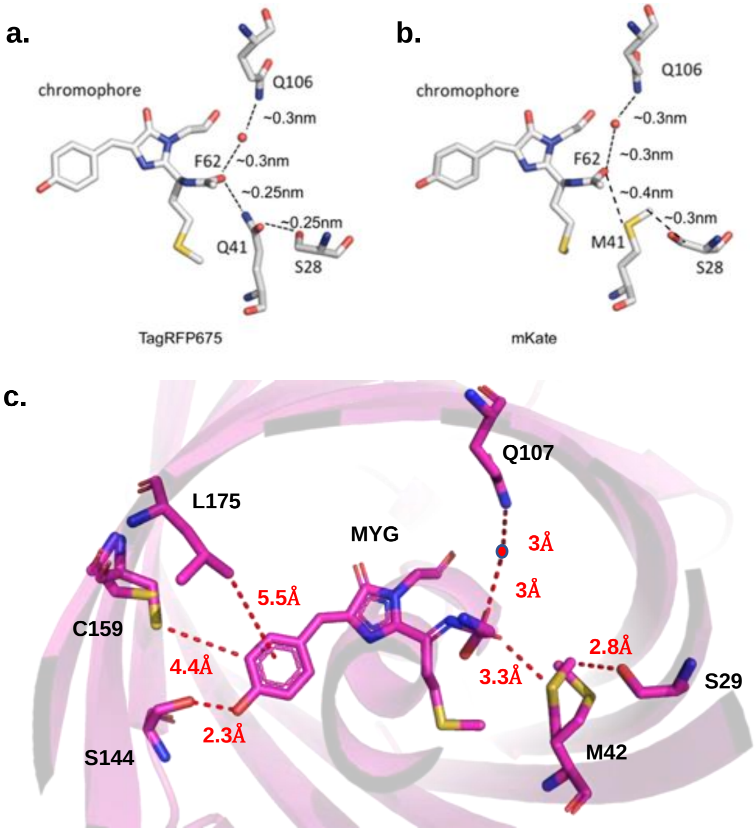


**Figure S1:** The hydrogen bonding network in (a) TagRFP-675 and (b) mKate. Images modified from Konold *et al*.[3] Distances were calculated using the crystal structures of TagRFP-675 (PDB ID: 4KGE) and mKate (PDB ID:3BXA). Konold and co-workers describe an extensive network involving the Q106, S28 and M/Q41 residues with a crystallized water molecule at the acylimine end of these RFPs (c) The crystal structure of FR (PDB ID: 6U1A) also reveals a similar arrangement. Relevant positions and the distances from the chromophore in the FR structure are shown.

### S4: Additional photophysical and biochemical characterization

***S4a. Excitation versus absorption spectra:*** Excitation traces and absorption spectra overlaid with each other for FR and the FR mutants generated in this study. In the main text, our analysis for absorption wavelengths, calculations of Stokes shift, etc., were done with respect to the absorption spectra for the relevant FPs.

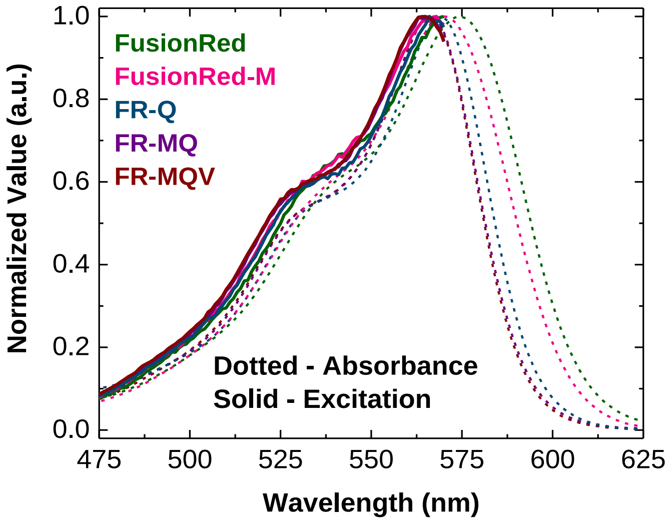


**Figure S2:** Overlaying the absorption and the excitation spectra for the FPs relevant to this study. The excitation spectra (solid) and absorption spectra (dotted). The y-axis indicates the normalized value of absorption/excitation in normalized arbitrary units (a.u.).

***S5b. Spectral characteristics:*** We observed that both the absorption and emission spectral width reduced as mutations were progressively incorporated into FR in the development of FR-MQV. Table S3 below summarizes the spectral changes that we observed in this series of mutants.

**Table S3:** Spectral characteristics of the FR family of proteins investigated in this study. There is a progressive blue shift in the series. The change in Stokes shift, unlike mKate-Q, is minimal for the M42Q mutants of FR. Integrated absorption and emission for peak-normalized spectra indicate the progressive narrowing of spectra in the series.

| Protein | $\lambda_{abs\_max}$ (nm) | Integrated absorption (a.u.) (400-650nm) | $\lambda_{em\_max}$ (nm) | Integrated Emission (a.u.) (400-650nm) | Stokes shift (nm) | Stokes shift ($cm^{-1}$) |
|---|---|---|---|---|---|---|
| FR-MQV | 566 | 58.3 | 585 | 51.7 | 19 | 574 |
| FR-MQ | 567 | 60.0 | 586 | 52.5 | 19 | 572 |
| FR-Q | 568 | 61.7 | 587 | 52.6 | 19 | 570 |
| FR-M | 571 | 66.0 | 590 | 62.2 | 19 | 595 |
| FR | 574 | 68.8 | 596 | 64.1 | 22 | 643 |

***S4c. Normalization with respect to the 278nm Trp peak indicates higher peak absorption for M42Q mutants of FR:*** FR and FR mutants with 42Q have the same number of Trp residues. Normalizing with respect to the absorption peak of tryptophan at ~280 nm should reflect the true nature of spectral width for the RFP peak. On doing so we observe the red emission peak is indeed narrower and shifted blue relative to the parent protein.

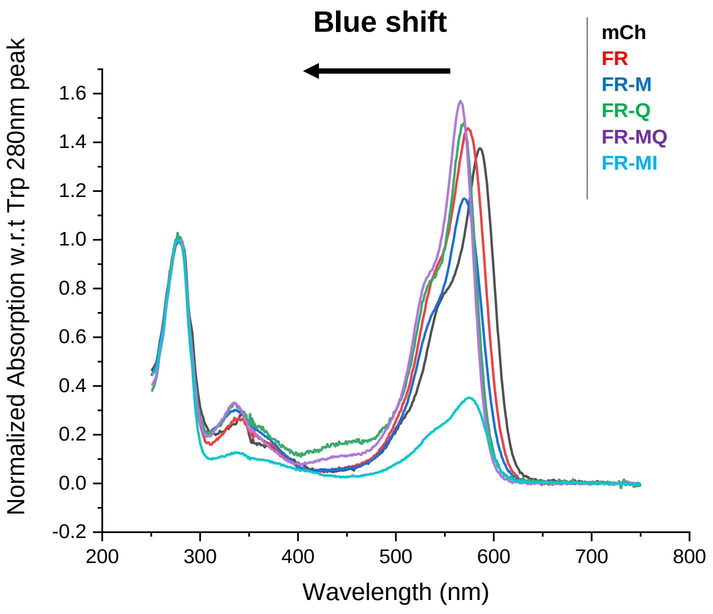


**Figure S3:** Normalizing the absorption spectra with respect to the Trp 280 nm feature reveals that M42Q changes the peak absorption. FR-MQ has a ~20% higher peak absorbance compared to FR.

***S4d. pH titrations and alkali denaturation:*** We collected absorption spectra for FPs *in vitro* at pH values 2–14. At acidic pHs FR-MQV resembles FR and FR-M, but at basic pH values, FR-MQV lacks the 450 nm degradation product that is present in FR-M and many other red FPs, such as mScarlet-I. Instead, there is a single product of alkali denaturation at 380 nm whenever the 42Q mutation is incorporated into the FR protein. However, this effect is not seen in mKate-Q, which, like the parent mKate and other RFPs, such as mScarlet-I (also mCherry, mScarlet, mRuby3; data not shown), degrades to only the 450 nm hydrolysis product.

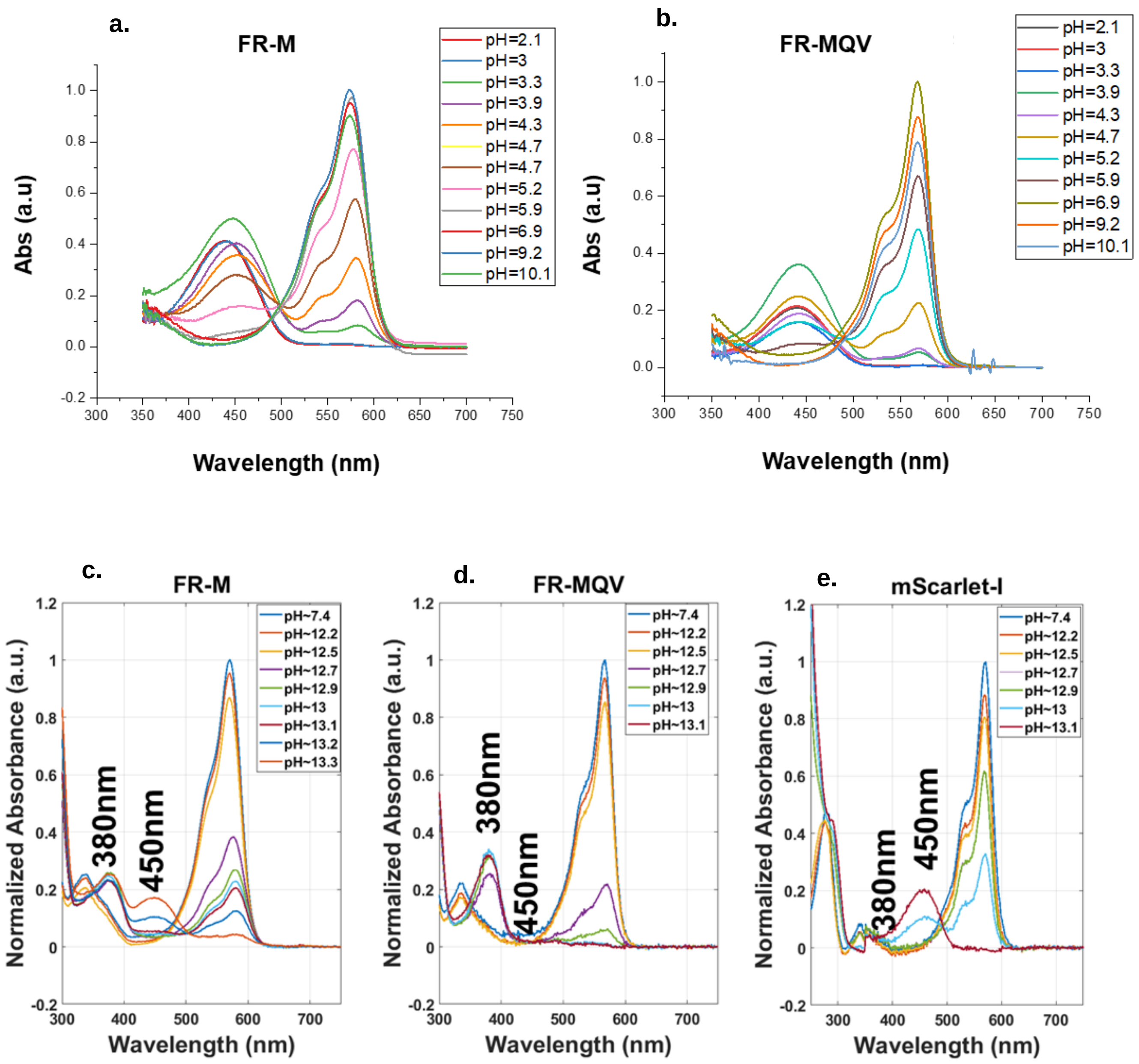

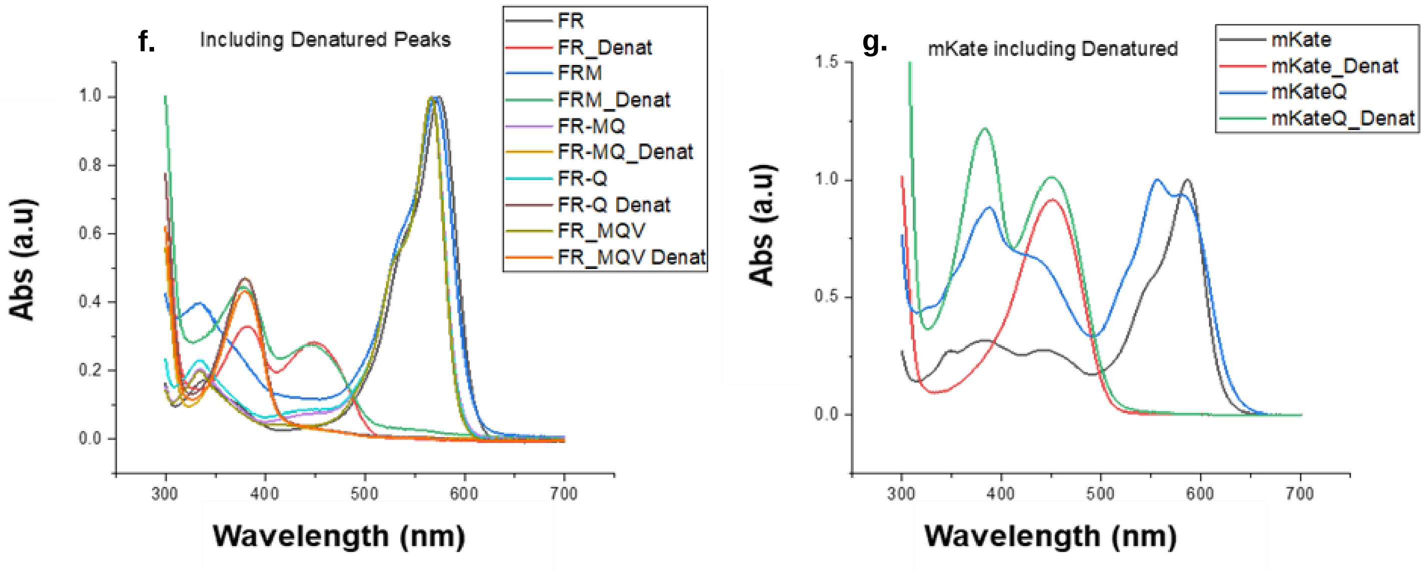


**Figure S4:** pH titrations from pH 2 to 10 of (a) FR-M and (b) FR-MQV show similar behavior for the functional form and the acid-degraded form of the FPs. Titration in the basic range (pH >12) shows that (c) FR-M breaks down into two products of alkali hydrolysis, with the 380 nm peak formed at a lower pH than the 450 nm peak; (d) FR-MQV displays only one product of alkali degradation at 380 nm and; (e) mScarlet-I (along with other RFPs) shows only the 450 nm peak for alkali degradation. (f) All constructs with the M42Q mutation behave similarly, but such an effect is not seen in (g) mKate-Q. The y-axis indicates the normalized value of absorption in normalized arbitrary units (a.u.).

***S4e. Green emission peak:*** The formation of some RFP chromophores is accompanied by the formation of a green chromophore that lacks the extension of the acylimine moiety[4]. This is undesirable as it decreases the amount of mature red species present in a protein sample. Protein engineering efforts, particularly those that involve mutations internal to the beta barrel often disrupt the chromophore maturation pathway and hence inadvertently result in an increase in the immature or green species. FR mutants developed in this work employed multiple internal mutations, emission spectra were collected in the 465–750 nm window, with excitation at 450 nm for Figure S5. It is evident that the M42Q mutation does not increase the green emission peak relative to FR (both show a shoulder ~20% of the major red peak) suggesting a properly mature red chromophore. However, the same mutation in mKate changes the spectrum considerably. Even the incorporation of a similar-sized, but aliphatic residue, isoleucine, produces a large green emission peak.

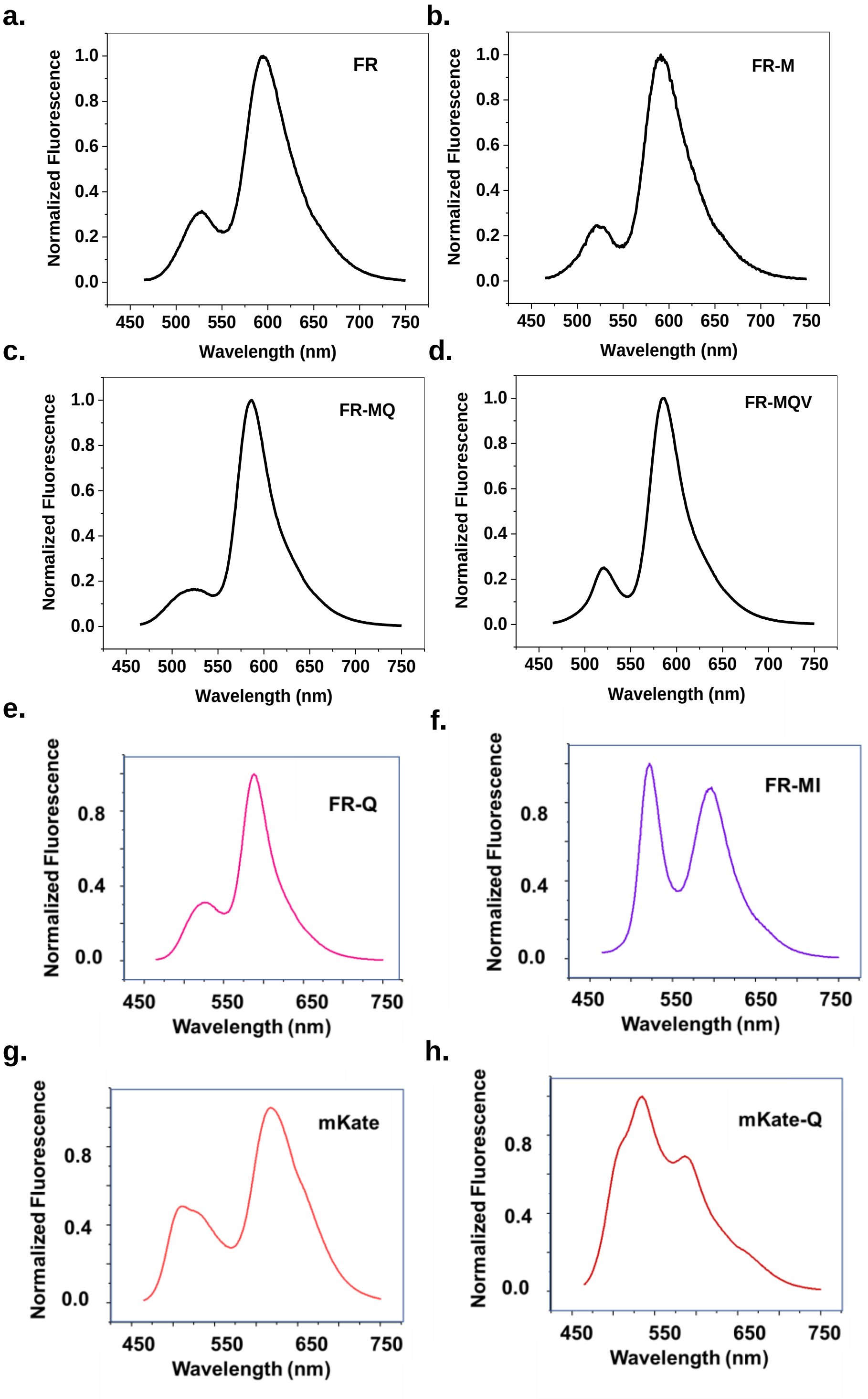


**Figure S5:** Emission spectra recorded with respect to excitation at 450 nm for (a) FR, (b) FR-M, (c) FR-MQ, (d) FR-MQV, (e) FR-Q, (f) FR-MI, (g) mKate, and (h) mKate-Q. FR mutants except FR-MI exhibit minimal green fluorescence, suggesting proper red chromophore maturation. mKate-Q also displays a significant green peak.

***S4f. Excitation-dependent emission:*** Konold and co-workers previously demonstrated the existence of multiple non-interconverting hydrogen-bonded conformations of TagRFP-675 and mKate-Q.[3] We excited the proteins in this study at 500 nm, 525 nm and 550 nm to test for different emission species in the red window as shown in Figure S6. mKate-Q displays multiple emission profiles with interchanging populations (peak height). However, there are no shifts for the parent mKate and other FR mutants.

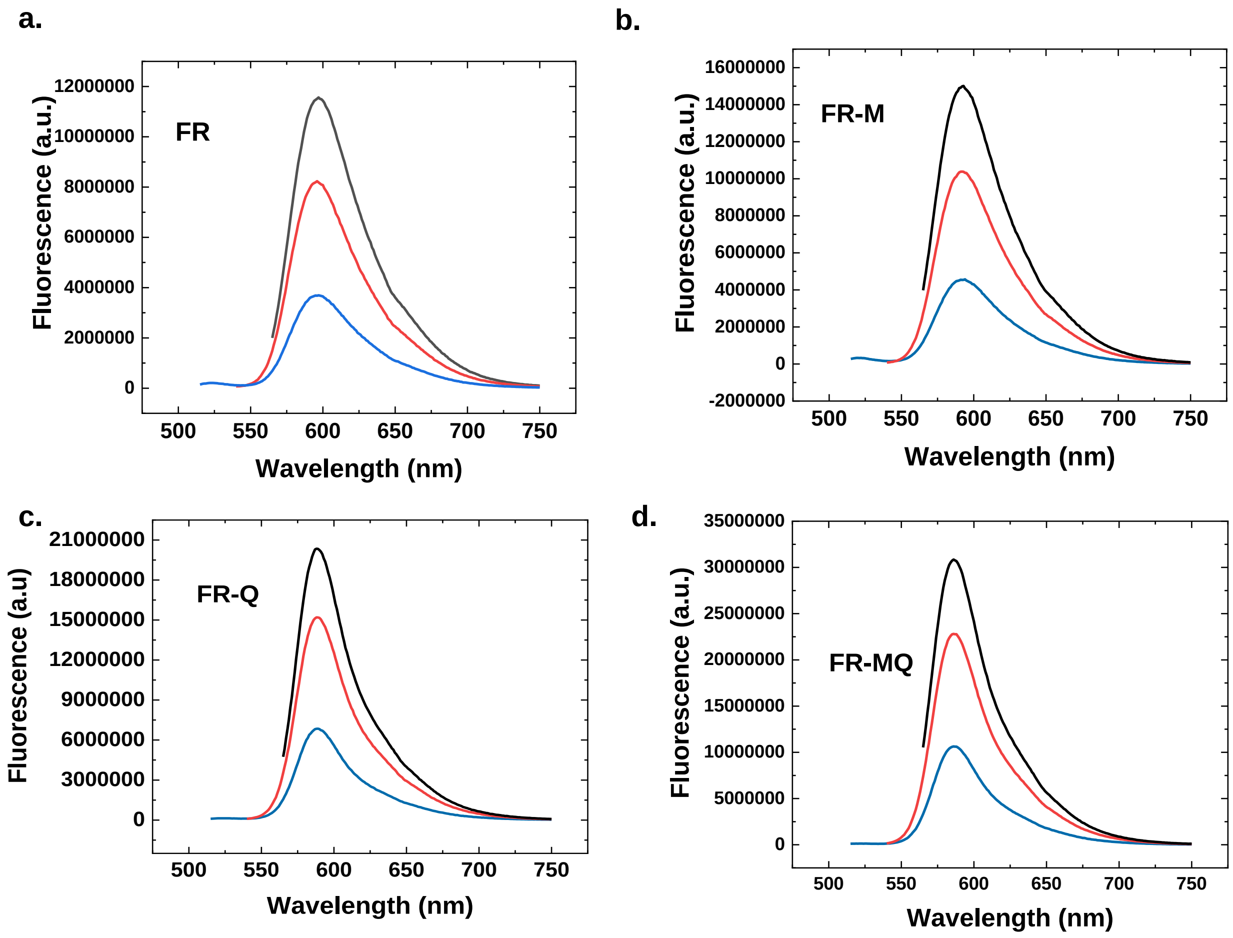

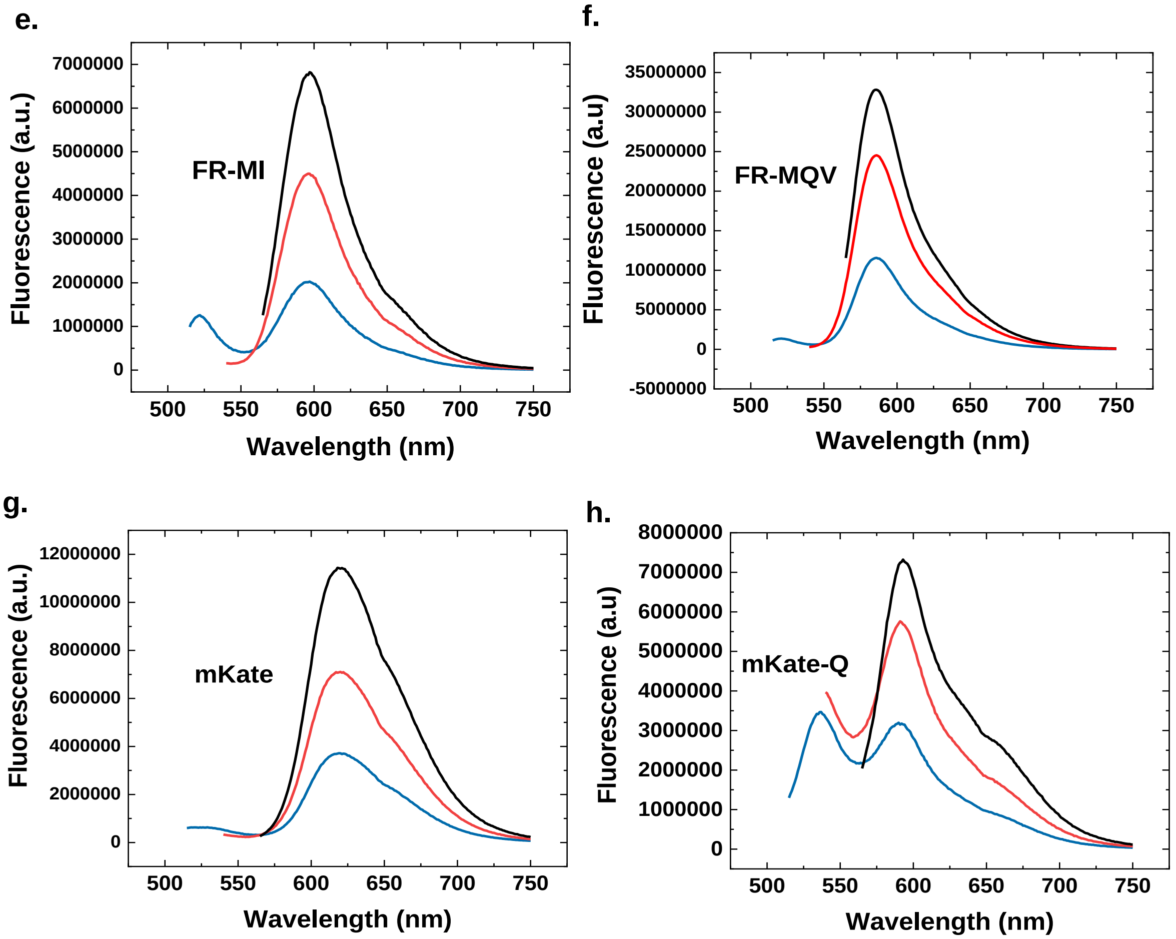


**Figure S6 :** Emission spectra of (a) FR, (b) FR-M, (c) FR-Q, (d) FR-MQ, (e) FR-MI, (f) FR-MQV, (g) mKate, and (h) mKate-Q, with excitation at 500 (blue), 525 (red) and 550 (black) nm. Consistent with previous results, mKate-Q has multiple emissive species.

***S4g. Lifetime measurements using time-correlated single photon counting (TCSPC):*** Fluorescence lifetime was measured using TCSPC by exciting with a 560 nm pulsed laser and collecting emission in two windows using an appropriate filter set. The changes in populations (amplitudes of fits in lifetime decay) and the changes in average lifetime indicate the existence of multiple emissive configurations. Data are presented in Table S4.

**Table S4:** TCSPC-based lifetime measurements using pure protein samples. Independent trials are separated by commas, where each **τ** indicates the component of lifetime in ns and the % intensity for that component is reported in brackets. As the L175M, C159V and M42Q mutations are added on FR, there is a progressive increment of **τ** in the major component of fluorescence decay. The mKate-42Q mutant displays a population shift based on the emission window suggesting multiple non-interconverting conformers and fits to a tri-exponential function. Other FPs do not show a strong dependence on emission windows, with similar populations (based on intensity % in the fits) and time constants fitting best to bi-exponential functions.

| Protein | Emission window (600 ± 30 nm) | | | | Emission window (670 ± 30 nm) | | | |
|---|---|---|---|---|---|---|---|---|
| | **$\tau_1$ (%Int)** | **$\tau_2$(%Int)** | **$\tau_3$(%Int)** | **$\tau_{av}$** | **$\tau_1$ (%Int)** | **$\tau_2$(%Int)** | **$\tau_3$(%Int)** | **$\tau_{av}$** |
| **FR** | 1.40(64), 1.44(63), 1.42(64) | 2.49(36), 2.51(37), 2.51(37) | - | 1.78±0.04 | 1.37(39) | 2.36(61) | - | 1.76 |
| **FR-M** | 1.55(38), 1.56(33), 1.49(33) | 2.43(62), 2.47(67), 2.43(67) | - | 2.13±0.05 | 1.70(53) | 2.54(47) | - | 2.09 |
| **FR-Q** | 1.50(55), 1.52(53), 1.56(52) | 2.92(45), 2.93(47), 2.97(48) | - | 2.10±0.05 | 2.93(44) | 1.51(56) | - | 2.13 |
| **FR-MQ** | 1.74(35), 1.87(40), 2.05(40) | 2.82(65), 2.91(60), 3.02(56) | - | 2.45±0.08 | 2.98(56) | 1.85(44) | - | 2.47 |
| **FR-MI** | 2.22(55), 2.21(54) | 1.21(45), 1.29(46) | - | 0.26±0.04 | 2.16(56) | 1.16(44) | - | 1.72 |
| **FR-MQV** | 3.23(68), 3.15(69), 3.14(69) | 2.03(32), 1.94(31), 2.01(31) | - | 2.77±0.07 | 3.13(73) | 1.91(27) | - | 2.80 |
| **mKate** | 2.40(92), 2.42(90), 2.37(93) | 0.87(8), 0.86(10), 0.80(8) | - | 2.26±0.07 | 2.38(92) | 8.21(8) | - | 2.25 |
| **mKate-Q** | 3.72(52), 3.61(48), 3.65(55) | 1.29(40), 1.25(41), 1.32(39) | 0.21(8), 0.19(9), 0.46(6) | 2.16±0.05 | 3.62(36) | 1.27(61) | 0.24(3) | 2.05 |
| **mCherry** | 1.63(88), 1.75(70) | 0.59(12), 1.06(30) | - | 1.67±0.07 | - | - | - | - |
| **mScarlet** | 3.82(98), 3.80(97), 3.93(99) | 1.27(2), 1.18(3), 1.31(1) | - | 3.87±0.07 | - | - | - | - |
| **mScarlet-I** | 3.41(92), 3.57(86), 3.51(88) | 0.92(8), 1.63(14), 1.51(12) | - | 3.26±0.07 | - | - | - | - |
| **mRuby3** | 2.79(95) | 1.16(5) | - | 2.71 | - | - | - | - |

***S4h. pKa measurements:*** Table S5 shows the pKa values of relevant mutants in this study. The pKa was obtained by taking three measurements for each pH data point and fitting the curve to a sigmoidal function (Figure S7). The half value of the maximum fluorescence was calculated to be the pKa of the protein. The traces for mScarlet, mScarlet-I and FR-MQV are shown in Figure S7. Both mScarlet and mScarlet-I scale well with the reported values by Bindels *et al.*[5]

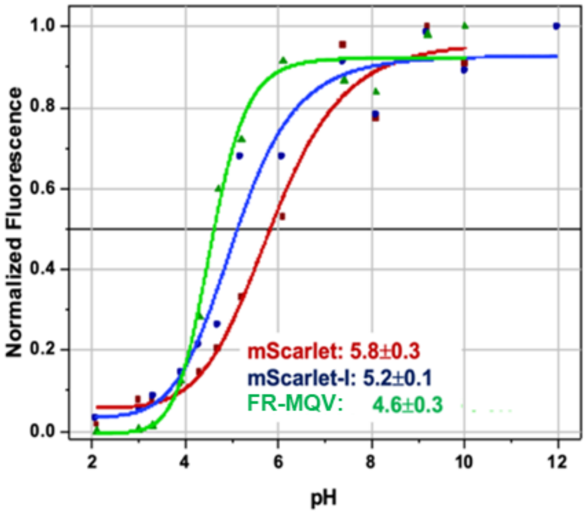


**Figure S7:** Plot of fluorescence versus pH for FR-MQV, mScarlet and mScarlet-I. The pKa is obtained from the 0.5 fractional fluorescence point. FR-MQV retains the low pKa value of the FR family of proteins.

**Table S5:** The pKa values measured in this study versus the values reported for known RFPs in the literature.

| Protein | pKa Calculated | pKa Reported |
|---|---|---|
| FR | 4.5±0.02 | 4.5[6,7] |
| FR-M | 4.7±0.03 | 4.8[6,7] |
| FR-Q | 4.3±0.02 | - |
| FR-MQ | 4.4±0.01 | - |
| FR-MQV | 4.6±0.02 | - |
| mScarlet | 5.8±0.03 | 5.3[5] |
| mScarlet-I | 5.2±0.01 | 5.4[5] |

***S4i. Bacterial brightness:*** The family of FR mutants relevant to this study were expressed in bacteria to test bacterial brightness in a droplet microfluidic screening platform (summarized in Figure S8).[8] Brightness seems to scale well with increasing molecular brightness with FR-MQV being ~2-fold brighter than FR. Details of the experiment are described in the methods and materials section of the main text of the article.

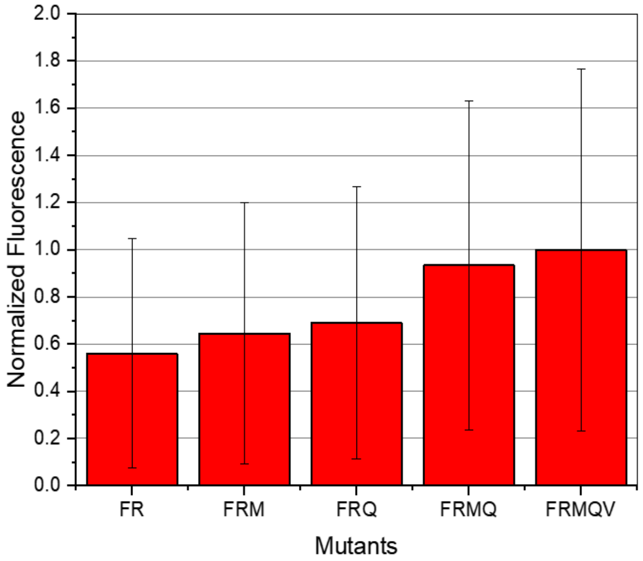


**Figure S8:** The brightness (normalized with respect to FR-MQV) recorded for the family of FR mutants relevant to this study. We see a progressive increase in bacterial brightness for the family, with FR-MQV being ~2-fold brighter than FR.

***S4j: Photobleaching experiments:*** The photobleaching traces of FR, FSD-9 (FR-C159V) and FSD-11 (FR-C159L) are shown in Figure S9 (a). The fast fluorescence decay process may result from transition to a dark state, as has been observed previously for FPs.[9] The incorporation of a small aliphatic group at position 159 in FSD-9 eliminates the tendency of the FP to undergo the dark state conversion process. Thus, the photobleaching profile changed from a bi-exponential-like decay in FR to a mono-exponential-like decay in FSD-9 and FSD-11. It is worth noting that FSD-11 shows a slow photoactivation process in ~50 s under the irradiance regime used in this study. The tendency for photo-switching, or reversible photobleaching, is the lowest for FR-C159V, followed by FR-C159L, and the highest in FR, suggesting that an aliphatic group at the 159 position may help to reduce the cis–trans isomerization of the chromophore.

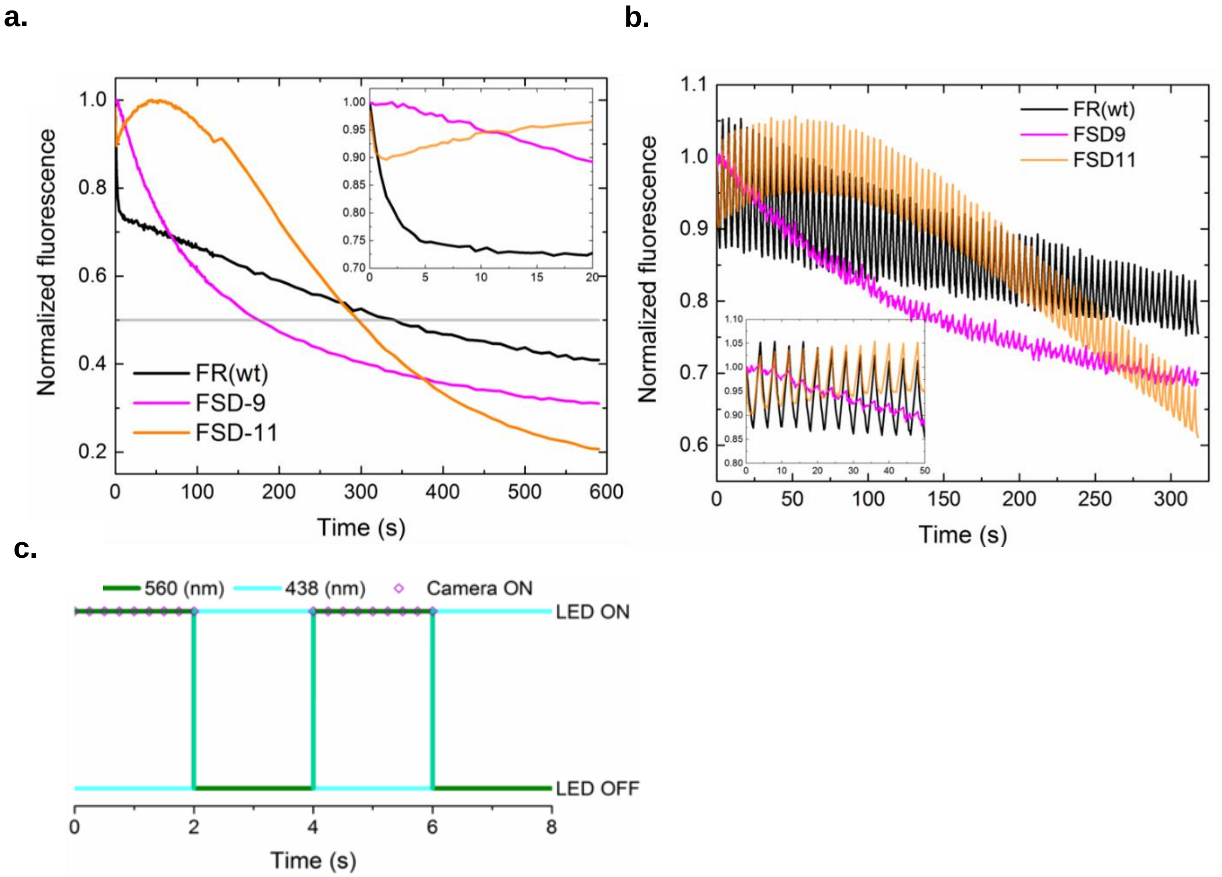


**Figure S9:** (a) FR, FSD-9 (FR-C159V) and FSD-11 (FR-C159L) expressed in bacteria were photobleached with 560 nm LED at a constant irradiance of ~4.9 W/cm$^2$. The inset exhibits the traces of the first 20 seconds. (b) Fluorescence signals under alternating illumination—each cell was illuminated with 560 nm LED for 2 s, then only 438 nm LED for another 2 s and then the illumination cycle was repeated. The camera only recorded signals when 560 nm LED was turned on. The inset shows the traces of the first 50 minutes. (c) The alternating LED light sources are illustrated for the first 8 s. The irradiance was ~4.9 W/cm$^2$ for 560 nm and ~4.5 W/cm$^2$ for 438 nm throughout the experiment. The camera recorded nine frames in 2 s when the 560 nm LED was turned ON.

**S5: Cellular Assays: brightness, maturation and cytotoxicity**

***S5a. Brightness assays:*** Brightness was measured using FACS and confocal microscopy. The Methods and Materials section of the main text describes the measurement protocols. Table S6 reports the values and number of biological replicates used for measuring the brightness through FACS. Each biological replicate had ~3 technical replicates of ~10000 HeLa cells each. Table S7 reports the number of HeLa cells analyzed in each sample dish and the mean intensity of each dish.

**Table S6:** Mean brightness from FACS measurements with standard deviation error from multiple biological replicates. Mean brightness measurements were normalized to FR.

| Protein | Biological Replicates | Mean Brightness |
|---|---|---|
| FR | 5 | 100±0 |
| FR-M | 4 | 191±31 |
| FR C159V | 2 | 164±17 |
| FR-M C159V | 2 | 266±31 |
| FR- Q | 2 | 214±18 |
| FR-MQ | 3 | 315±43 |
| FR-MQV | 4 | 509±20 |
| mCherry | 5 | 176±39 |
| mScarlet | 4 | 715±51 |
| mScarlet-I | 1 | 355 |

**Table S7:** Mean brightness from confocal microscopy measurements with standard deviation error from the number of cells indicated in each dish. Mean intensity measurements were normalized to FR.

| Protein | # Cells | Mean Intensity (x100) |
|---|---|---|
| mCherry-1 | 51 | 77±55 |
| mCherry-2 | 38 | 171±137 |
| FR-1 | 12 | 100±68 |
| FR-2 | 61 | 100±78 |
| mScarlet-1 | 131 | 250±194 |
| mScarlet-2 | 71 | 379±273 |
| FR-MQV-1 | 62 | 231±154 |
| FR-MQV-2 | 48 | 215±187 |
| Untransfected-1 | 513 | 4±2 |
| Untransfected-2 | 469 | 5±3 |

***S5b. Cytotoxicity assay*:** The detailed protocol for this assay is reported in the Materials and Methods section. Briefly, mammalian cells expressing either EGFP or one of the RFP clones were mixed in a 50:50 ratio by volume. Initially, cells expressing EGFP and each RFP were FACS screened individually (~5000 cells) as controls. Part of mixture was analyzed by FACS to quantify the number of cells carrying EGFP and the RFP at the start (Day 2). The remaining mixture was re-plated. The screens for the re-plated mixtures were repeated after another 4 days of growth (Day 6). Ratios of RFP: EGFP mixtures were calculated for technical replicates for Day 2 and Day 6. The change in the ratio of RFP to EGFP for each replicate is a measure of the relative cytotoxicity of the RFP to EGFP. Figure S10 is a graphical representation of the assay. We found that FR and FR-MQV are reproducibly less cytotoxic than EGFP. In contrast, mCherry was consistently more cytotoxic than EGFP, whereas for mScarlet, in one case the RFP was more cytotoxic than EGFP and in another case it was less cytotoxic.

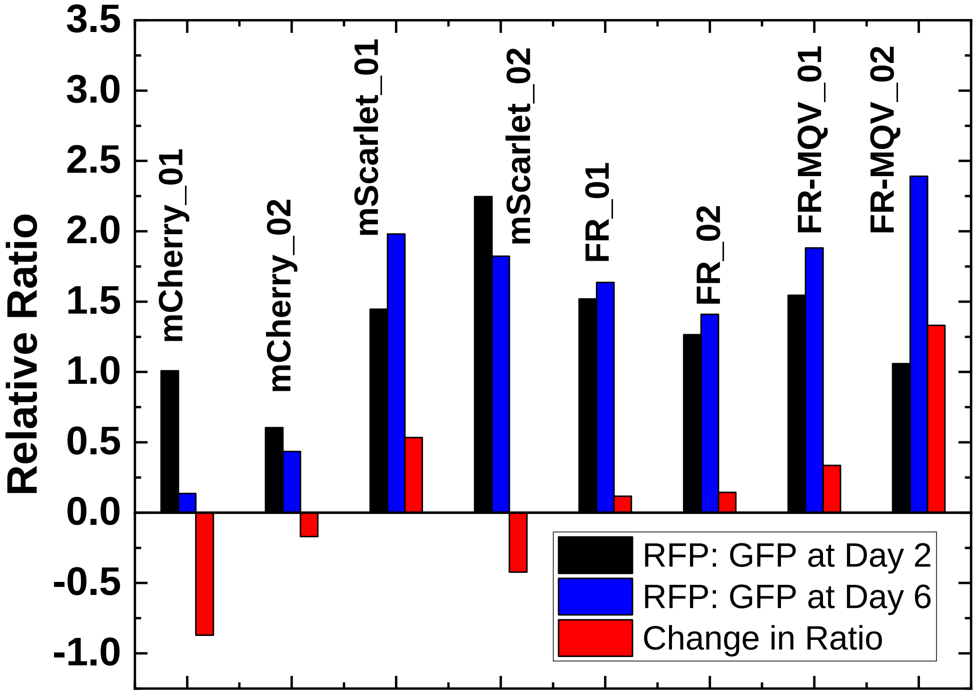


**Figure S10:** Cytotoxicity assay: The black bars represent the RFP:EGFP ratio in a mix on Day 2. The bars represent the RFP:EGFP ratios measured on Day 6. On each day ~10,000 cells were measured using FACS. The change in the ratio is represented as red bars. FR and FR-MQV are consistently less cytotoxic than EGFP.

**S5c. Chromophore maturation kinetics:** Details of maturation kinetics are provided in the Methods and Materials section. Briefly, FPs were expressed in *E. coli*. After induction of protein expression, cultures were treated with chloramphenicol to halt new protein production and both the fluorescence and optical density were measured over time. An increase in fluorescence (after normalization to optical density) indicates an increase in chromophore formation. FR and FR-MQV show similar maturation kinetics. The mScarlet and mScarlet-I values are comparable to those reported in the literature. Hence, the mutations in FR-MQV do not appear to perturb maturation and folding of the FP at 37ºC (temperatures used for mammalian cell growth/imaging experiments).

**a.** **b.**

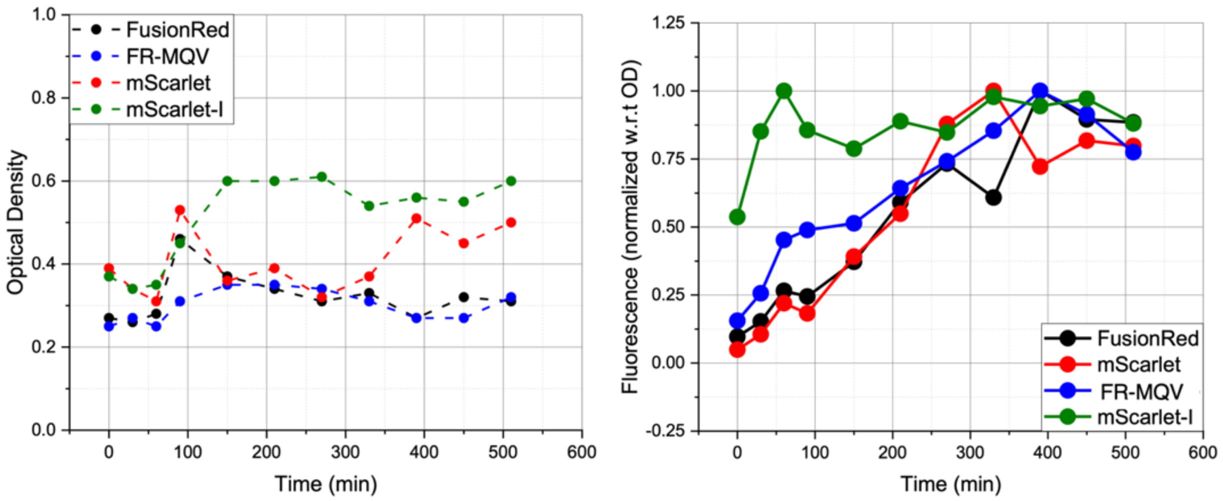


**Figure S11:** (a) The optical density (OD) of bacterial cultures over time after addition of chloramphenicol at t = 0. The ODs remained fairly constant, suggesting the action of chloramphenicol stalling the growth of bacterial cells in the log phase. (b) The measured fluorescence normalized and scaled with respect to the observed ODs at each time point.

**Table S8:** Measured versus the reported maturation times ($t_{50}$) for the RFPs investigated in this study.

| Fluorescent Protein | ~ $t_{50}$ 37ºC | Reported $t_{50}$ (min) |
|---|---|---|
| mScarlet-I | 45 min | 36[5] , 25[10] |
| mScarlet | 165 min | 174[5], 132 [10] |
| FR | 195 min | 130[6] |
| FR-MQV | 195 min | - |

**S6: Extinction coefficient and quantum yield estimation:**

***S6a. Extinction coefficient calculation based on SDS-PAGE:***

*Calculation of extinction coefficient was based on the original FusionRed[6] article for FPs exhibiting backbone cleavage:*

$$\epsilon_{RFP} = \frac{Abs_{max\ RFP}}{\left(\frac{Abs_{380nm}}{\epsilon_{380nm}}\right) + \left(\frac{Abs_{450nm}}{\epsilon_{450nm}}\right)}$$

Where $Abs_{max\ RFP}$ is the maximum absorbance for the RFP in the undenatured absorption spectrum and $Abs_{380nm}$ and $Abs_{450nm}$ were the values of absorbance at 380 nm and 450 nm in the alkali denatured spectrum. We used previously reported values of $\epsilon_{380nm}$= 70,500 $M^{-1}cm^{-1}$ and $\epsilon_{450nm}$= 44,000 $M^{-1}cm^{-1}$.[6,7] SDS-PAGE was used to assess the purity of proteins extracted from *E. coli* and to examine whether FR-MQV exhibited backbone cleavage like the parent FR such that the mathematical formula presented above could be used to calculate the extinction coefficient of the RFP. Proteins were loaded at 1X and 2X (~10 μM) concentrations. Cytochrome-C (Cyt-C) was included as an additional size marker and EGFP was included as a negative control uncleaved FP. The results for FR are consistent with previously reported data, therefore we used the above-mentioned relationship to estimate the extinction coefficient of FR and its family of mutants.

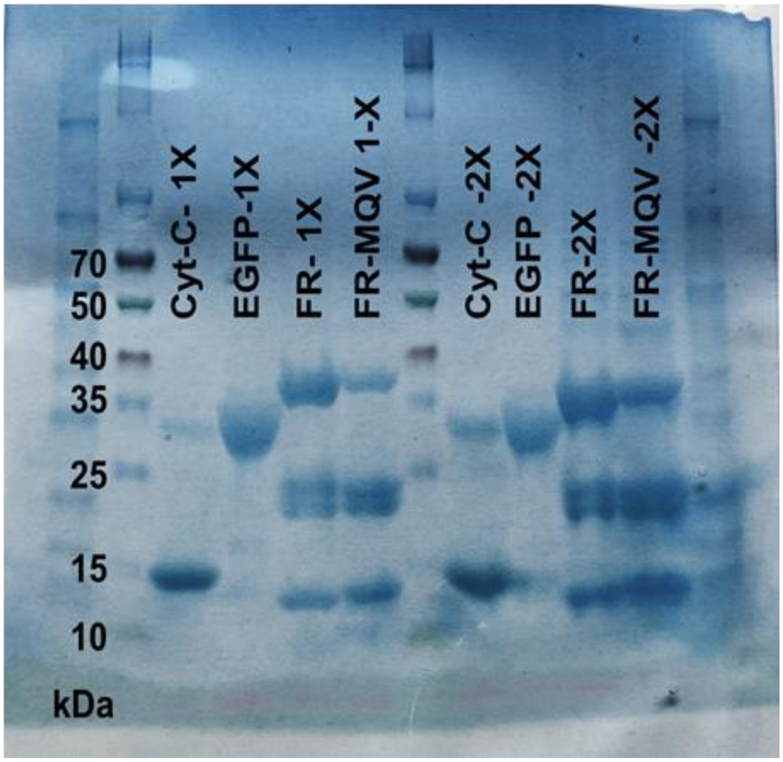


**Figure S12:** Gradient (4-20%) SDS-PAGE of FR and FR-MQV with appropriate controls shows backbone cleavage for FR-MQV.

***S6b. Quantum yield measurements:*** The quantum yield (QY) for purified protein samples was measured as described in the Methods and Materials. The data follows expected linear trends as shown in Figure S13. Table S9 presents the number of independent trials and the standard deviation error observed in measurement for each FP.

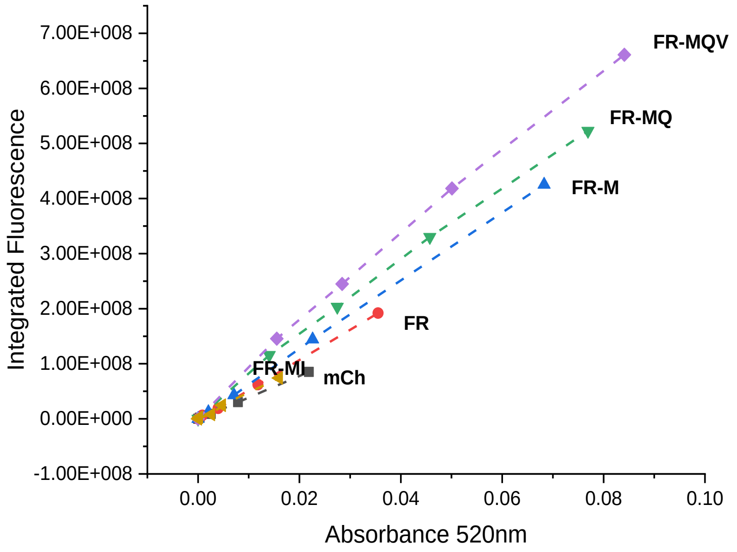


**Figure S13:** The linear trend of higher integrated fluorescence against absorption measurements under serial dilution for one of the independent measurements. Higher slope values indicate higher quantum yields.

**Table S9:** Measured mean QY and standard deviation errors for FPs in this study. QY measurements are susceptible to random and systematic errors. Cranfill *et al.* described protocols to minimize such errors.[11] Steps such as pH control, minimizing temperature fluctuations, gentle thawing of FPs and usage of fresh protein were taken into consideration. Replicate measurements for QY of relevant proteins were taken from batches of FPs prepared through independent transformations, growth and purification protocols. Cresyl Violet, mCherry and mScarlet were usually used as multiple references for the measurements.

| **Protein** | **Mean QY** | **Trials** |
|---|---|---|
| FR-MI | 26±4 | 3 |
| FR-MQ | 43±3 | 4 |
| FR-MQV | 53±3 | 4 |
| FR-Q | 33±1 | 4 |
| FR | 24±4 | 4 |
| FR-M | 34±2 | 4 |
| mCherry | 22 (ref) | 4 |
| mKate | 33 | 1 |
| mKate-42Q | 17 | 1 |
| mRuby3 | 43 | 1 |
| mScarlet | 72±4 | 3 |
| mScarlet-I | 59±2 | 3 |

#### S7: FR Evolution Table:

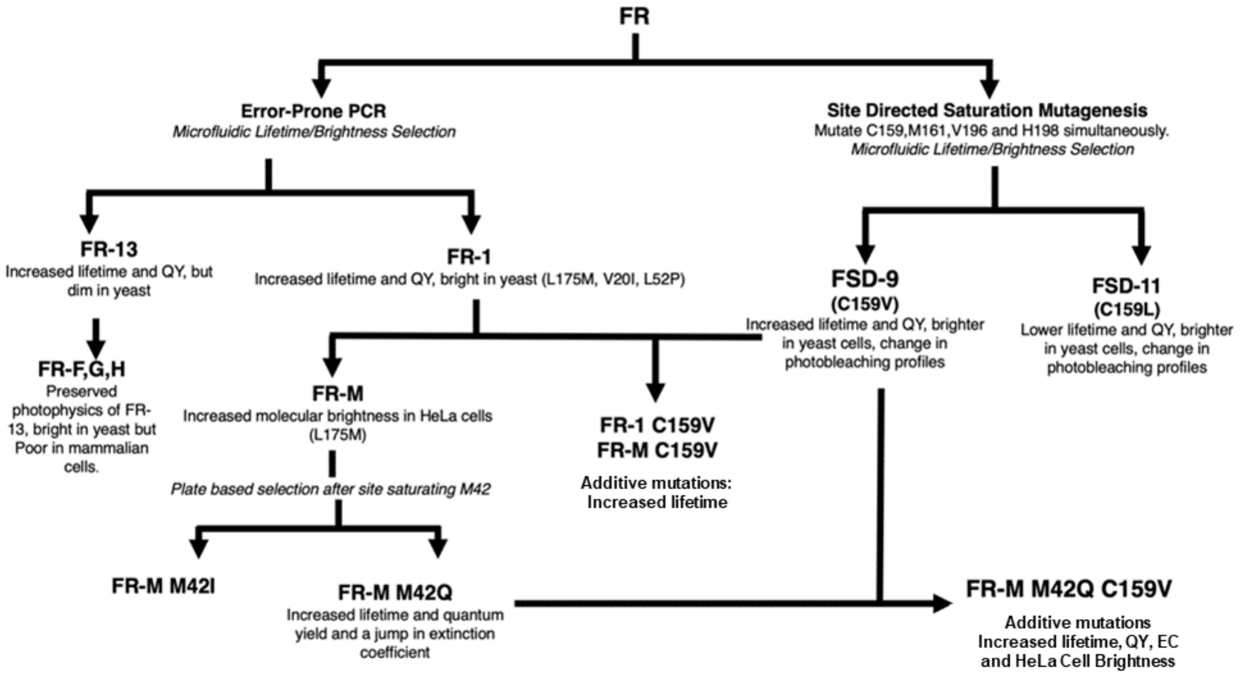


**Figure S14:** The evolution of the FR family of proteins. The pathways indicate the various engineering strategies that led to the development of FR-MQV.[7] All amino acid positions have been numbered with respect to the parent FR numbering as per Table S1.
